# Global organization of neuronal activity only requires unstructured local connectivity

**DOI:** 10.1101/2020.07.15.205013

**Authors:** David Dahmen, Moritz Layer, Lukas Deutz, Paulina Anna Dąbrowska, Nicole Voges, Michael von Papen, Thomas Brochier, Alexa Riehle, Markus Diesmann, Sonja Grün, Moritz Helias

**Affiliations:** Institute of Neuroscience and Medicine (INM-6) and Institute for Advanced Simulation (IAS-6) and JARA Institut Brain Structure-Function Relationships (INM-10), Jülich Research Centre, Jülich, Germany; RWTH Aachen University, Aachen, Germany; School of Computing, University of Leeds, Leeds, West Yorkshire, United Kingdom; Institut de Neurosciences de la Timone (INT), CNRS - Aix-Marseille University, Marseille, France; Department of Physics, Faculty 1, RWTH Aachen University, Aachen, Germany; Department of Psychiatry, Psychotherapy and Psychosomatics, School of Medicine, RWTH Aachen University, Aachen, Germany; Theoretical Systems Neurobiology, RWTH Aachen University, Aachen, Germany

## Abstract

Modern electrophysiological recordings simultaneously capture single-unit spiking activities of hundreds of neurons spread across large cortical distances. Yet this massively parallel activity is often confined to relatively low-dimensional manifolds. This implies strong coordination also among neurons that are most likely not even connected. Here, we combine in vivo recordings with network models and theory to characterize the nature of mesoscopic coordination patterns in macaque motor cortex and to expose their origin: We find that heterogeneity in local connectivity supports network states with complex long-range cooperation between neurons that arises from multi-synaptic, short-range connections. Our theory explains the experimentally observed spatial organization of covariances in resting state recordings as well as the behaviorally related modulation of covariance patterns during a reach-to-grasp task. The ubiquity of heterogeneity in local cortical circuits suggests that the brain uses the described mechanism to flexibly adapt neuronal coordination to momentary demands.

## Introduction

Complex brain functions require coordination between large numbers of neurons. Unraveling mechanisms of neuronal coordination is therefore a core ingredient towards answering the long-standing question of how neuronal activity represents information. Population coding is one classical paradigm (*1*) in which entire populations of neurons behave coherently, thus leading to positive correlations among their members. The emergence and dynamical control of such population-averaged correlations has been studied intensely (*2, 3, 4, 5*). More recently, evidence accumulated that neuronal activity evolves within manifolds, which implies even more involved ways of neuronal activity coordination (*6, 7, 8*): A small number of population-wide activity patterns, the neural modes, are thought to explain most variability of neuronal activity. In this case, individual neurons do not necessarily follow a stereotypical activity pattern that is identical across all neurons contributing to a representation. Instead the coordination among the members is determined by more complex relations. Simulations of recurrent network models indeed indicate that networks trained to perform a realistic task exhibit activity organized in manifolds (*9*). The dimensionality of such manifolds is determined by the structure of correlations (*10, 11*) and tightly linked to the complexity of the task the network has to perform (*12*) as well as to the dimensionality of the stimulus (*13*). Recent work has started to decipher how neural modes and the dimensionality of activity are shaped by features of network connectivity, such as heterogeneity of connections (*14, 15*), block structure (*16, 17*), and low-rank perturbations (*18*) of connectivity matrices, as well as connectivity motifs (*19, 20, 21*). Yet, these works neglected the spatial organization of network connectivity (*22*) that becomes more and more important with current experimental techniques that allow the simultaneous recording of ever more neurons. How distant neurons that are likely not connected can still be strongly coordinated to participate in the same neural mode is a widely open question.

To answer this question, we combine analysis of massively parallel spiking data from macaque motor cortex with the analytical investigation of a spatially organized neuronal network model. We here quantify coordination by pairwise covariances, which measure how temporal departures of the neurons’ activities away from their mean firing rate are correlated. We show that, even with only unstructured and short-range connections, strong covariances across distances of several millimeters emerge naturally in balanced networks if their dynamical state is close to an instability within a “critical regime”. While mean covariances are readily accessible by mean-field techniques and have been shown to be small in balanced networks (*3, 23*), explaining covariances on the level of individual pairs requires methods from statistical physics of disordered systems. With such a theory, here derived for spatially organized excitatory-inhibitory networks, we show that large individual covariances arise at all distances if the network is close to the critical point. These predictions are confirmed by recordings of macaque motor cortex activity. The long-range coordination found in this study is not merely determined by the anatomical connectivity, but depends substantially on the network state, which is characterized by the individual neurons’ mean firing rates. This allows the network to adjust the neuronal coordination pattern in a dynamic fashion, which we demonstrate through simulations and by comparing two behavioral epochs of a reach-to-grasp experiment.

## Results

### Macaque motor cortex shows long-range coordination patterns

We first analyze data from motor cortex of macaques during rest, recorded with 4 × 4 mm^2^, 100-electrode Utah arrays with 400 *µ*m inter-electrode distance (Figure 1*A*). The resting condition of motor cortex in monkeys is ideal to assess intrinsic coordination between neurons during ongoing activity. In particular, our analyses focus on true resting state data, devoid of movement-related transients in neuronal firing (see Methods). Massively-parallel single-unit spiking activity of ≈ 130 neurons per recording session, sorted into putative excitatory and inhibitory cells, shows strong spike-count covariances across the entire Utah array, well beyond the typical scale of the underlying short-range connectivity profiles (Figure 1*B*,*D*). Positive and negative covariances form patterns in space that are furthermore seemingly unrelated to the neuron types. All populations show a large dispersion of both positive and negative covariance values (Figure 1*C*). Given the classical view on pairwise covariances in balanced networks (*2, 24, 25, 23, 4*), this result comes unexpected: A common finding is that covariances averaged across many pairs of cells are small if the network dynamics is stabilized by an excess of inhibitory feedback; dynamics known as the ‘balanced state’ arise (*26, 27, 28*): Negative feedback counteracts any coherent increase or decrease of the population-averaged activity, preventing the neurons from fluctuating in unison (*23*). Breaking this balance in different ways leads to large covariances (*5, 29, 30*). Can the observation of significant covariances between individual cells across large distances be reconciled with the balanced state? In the following, we provide a mechanistic explanation.

**Figure 1:**
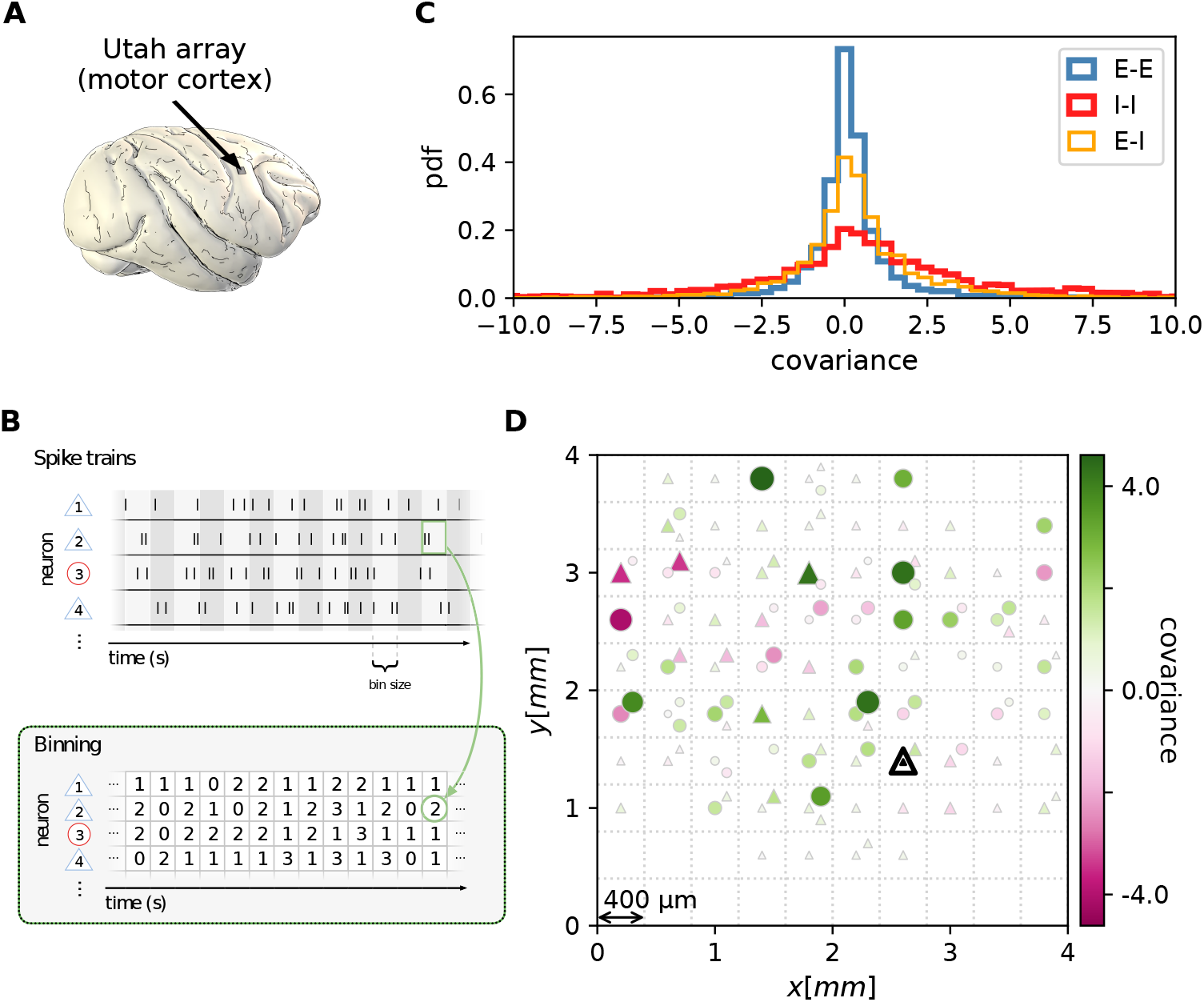
Salt-and-pepper structure of covariances in motor cortex. (*A*) Sketch of 10 × 10 Utah electrode array recording in motor cortex of macaque monkey during rest. (*B*) Spikes are sorted into putative excitatory (blue triangles) and inhibitory (red circles) single units according to widths of spike waveforms (see Appendix 1). Resulting spike trains are binned in 1 s bins to obtain spike counts. (*C*) Population-resolved distribution of pairwise spike-count covariances in session E2 (E-E: excitatory-excitatory, E-I: excitatory-inhibitory, I-I: inhibitory-inhibitory). (*D*)Pairwise spike-count covariances with respect to the neuron marked by black triangle in one recording (session E2, see Methods). Grid indicates electrodes of a Utah array, triangles and circles correspond to putative excitatory and inhibitory neurons, respectively. Size as well as color of markers represent covariance. Neurons within the same square were recorded on the same electrode. The source data and code used to create this figure are provided as fig1.zip.

### Multi-synaptic connections determine covariances

Connections mediate interactions between neurons. Many studies therefore directly relate connectivity and co-variances (*24, 25, 31, 32, 33*). From direct connectivity, one would expect positive covariances between excitatory neurons and negative covariances between inhibitory neurons and a mix of negative and positive covariances only for excitatory-inhibitory pairs. Likewise, a shared input from inside or outside the network only imposes positive covariances between any two neurons (Figure 2*A*). The observations that excitatory neurons may have negative covariances (Figure 1*D*), as well as the broad distribution of covariances covering both positive and negative values (Figure 1*C*), are not compatible with this view. In fact, the sign of covariances appears to be independent of the neuron types. So how do negative covariances between excitatory neurons arise?

**Figure 2:**
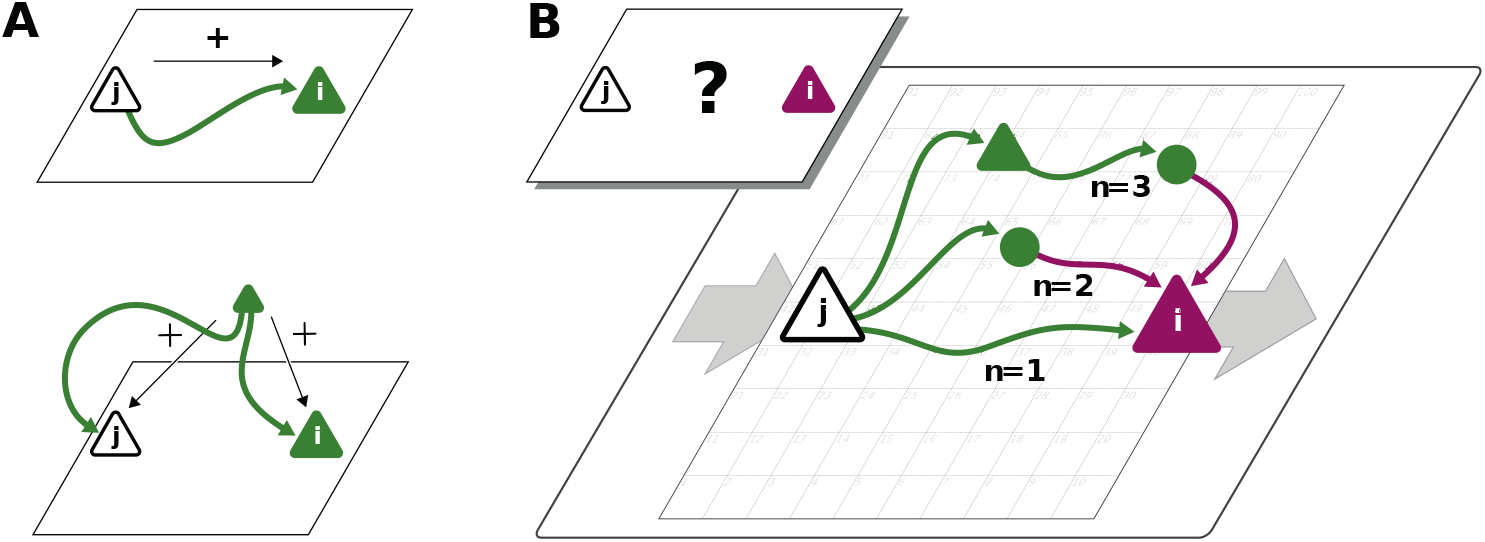
Covariances from direct and indirect connections. (*A*) Positive covariance (green neuron *i*) follows from direct excitatory connection (top) or shared input (middle). (*B*) Negative covariance (magenta) between two excitatory neurons cannot be explained by direct connections: Neuronal interactions are not only mediated via direct connections (*n* = 1; sign uniquely determined by presynaptic neuron type) but also via indirect paths of different length *n* > 1. The latter may have any sign (positive: green; negative: purple) due to intermediate neurons of arbitrary type (triangle: excitatory, circle: inhibitory).

The view that equates connectivity with correlation implicitly assumes that the effect of a single synapse on the receiving neuron is weak. This view, however, regards each synapse in isolation. Could there be states in the network where, collectively, many weak synapses cooperate, as perhaps required to form neuronal manifolds? In such a state, interactions may not only be mediated via direct connections but also via indirect paths through the network (Figure 2B). Such effective multi-synaptic connections may explain our observation that far apart neurons that are basically unconnected display considerable correlation of arbitrary sign.

Let us here illustrate the ideas first and corroborate them in subsequent sections. Direct connections yield covariances of a predefined sign, leading to covariance distributions with multiple peaks, e.g. a positive peak for excitatory neurons that are connected and a peak at zero for neurons that are not connected. Multi-synaptic paths, however, involve both excitatory and inhibitory intermediate neurons, which contribute to the interaction with different signs (Figure 2*B*). Hence, a single indirect path can contribute to the total interaction with arbitrary sign (*24*). If indirect paths dominate the interaction between two neurons, the sign of the resulting covariance becomes independent of their type. Given that the connecting paths in the network are different for any two neurons, the resulting covariances can fall in a wide range of both positive and negative values, giving rise to the broad distributions for all combinations of neuron types in Figure 1*C*. This provides a hypothesis why there may be no qualitative difference between the distribution of covariances for excitatory and inhibitory neurons. In fact, their widths are similar and their mean is close to zero (see Methods for exact values); the latter being the hallmark of the negative feedback that characterizes the balanced state. The subsequent model-based analysis will substantiate this idea and show that it also holds for networks with spatially organized heterogeneous connectivity.

To play this hypothesis further, an important consequence of the dominance of multi-synaptic connections could be that covariances are not restricted to the spatial range of direct connectivity. Through interactions via indirect paths the reach of a single neuron could effectively be increased. But the details of the spatial profile of the covariances in principle could be highly complex as it depends on the interplay of two antagonistic effects: On the one hand, signal propagation becomes weaker with distance, as the signal has to pass several synaptic connections. Along these paths mean firing rates of neurons are typically diverse, and so are their signal transmission properties (*34*). On the other hand, the number of contributing indirect paths between any pair of neurons proliferates with their distance. With single neurons typically projecting to thousands of other neurons in cortex, this leads to involved combinatorics; intuition here ceases to provide a sensible hypothesis on what is the effective spatial profile and range of coordination between neurons. Also it is unclear which parameters these coordination patterns depend on. The model-driven and analytical approach of the next section will provide such a hypothesis.

### Networks close to instability show shallow exponential decay of covariances

To gain an understanding of the spatial features of intrinsically generated covariances in balanced critical networks, we model a network of excitatory and inhibitory neurons on a two-dimensional sheet (Figure 3*A*, for details see Methods). The probability of two neurons being connected decays with distance on a characteristic length scale *d*. Previous studies have used linear response theory in combination with methods from statistical physics and field theory to gain analytic insights into both mean covariances (*2, 35, 24, 23*) and the width of the distribution of covariances (*15*). Field-theoretic approaches thereby were, however, restricted to purely random networks devoid of any network structure and thus not suitable to study spatial features of covariances. To analytically quantify the relation between the spatial ranges of covariances and connections, we therefore here develop a theory for spatially organized random networks with multiple populations. The model captures the predominant type of heterogeneity in cortical networks, namely the sparseness of connections.

**Figure 3:**
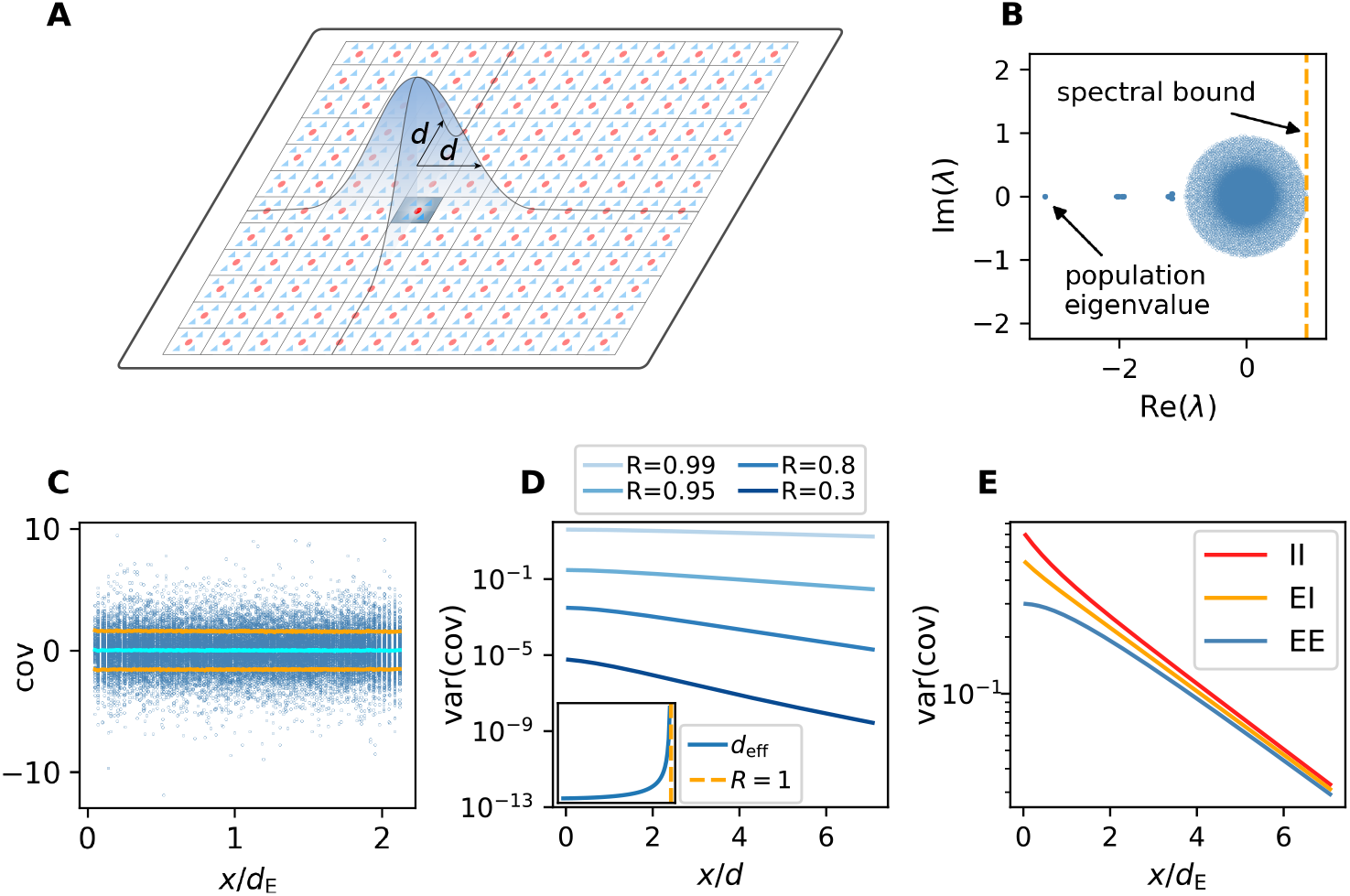
Spatially organized E-I network model. (*A*) Network model: space is divided into cells with four excitatory (triangles) and one inhibitory (circle) neuron each. Distance-dependent connection probabilities (shaded areas) are defined with respect to cell locations. (*B*) Eigenvalues *λ* of effective connectivity matrix for network in dynamically balanced critical state. Each dot shows the real part Re(*λ*) and imaginary part Im(*λ*) of one complex eigenvalue. The spectral bound (dashed vertical line) denotes the right-most edge of the eigenvalue spectrum. (*C*) Simulation: covariances of excitatory neurons over distance *x* between cells (blue dots: individual pairs; cyan: mean; orange: standard deviation; sample of 150 covariances at each of 200 chosen distances). (*D*) Theory: Variance of covariance distribution as a function of distance *x* for different spectral bounds of the effective connectivity matrix. *Inset*: effective decay constant of variances diverges as the spectral bound approaches one. (*E*) For large spectral bounds, the variances of EE, EI, and II covariances decay on a similar length scale. Spectral bound *R* = 0.95. Other parameters see Appendix 1-table 2. The source data and code used to create this figure are provided as fig3.zip.

A distance-resolved histogram of the covariances in the spatially organized E-I network shows that the mean covariance is close to zero but the width or variance of the covariance distribution stays large, even for large distances (Figure 3*C*). Analytically, we derive that, despite the complexity of the various indirect interactions, both the mean and the variance of covariances follow simple exponential laws in the long-distance limit (see Appendix 1 sections 3-11). These laws are universal in that they do not depend on details of the spatial profile of connections. Our theory shows that the associated length scales are strikingly different for means and variances of covariances. They each depend on the reach of direct connections and on specific eigenvalues of the effective connectivity matrix. These eigenvalues summarize various aspects of network connectivity and signal transmission into a single number: Each eigenvalue belongs to a “mode”, a combination of neurons that act collaboratively, rather than independently, coordinating neuronal activity within a one-dimensional subspace. To start with, there are as many such subspaces as there are neurons. But if the spectral bound in Figure 3B is close to one, only a relatively small fraction of them, namely those close to the spectral bound, dominate the dynamics; the dynamics is then effectively low-dimensional and multi-synaptic connections may become effective.

To quantify this idea, for the me<u>a</u>n covariance 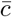we find that the dominant behavior is an exponential decay 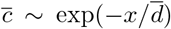 on a length scale 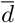 that is determined by the population eigenvalue (Figure 3*B*). Its position depends on the ratio between excitation and inhibition in the network and becomes more negative in more strongly inhibition-dominated networks. We show in Appendix 1 section 8.4 that this leads to a steep decay of mean covariances with distance. The variance of covariances, however, predominantly decays exponentially on a length scale *d*_eff_ that is determined by the spectral bound *R*, the largest real part among all eigenvalues (Figure 3*B, D*). In inhibition-dominated networks, *R* is determined by the heterogeneity of connections. For *R* ;S 1 we obtain the effective length scale

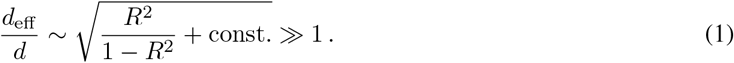

What this means is that precisely at the point where *R* is close to one, when the low-dimensional manifold forms, the length scale *d*_eff_ on which covariances decay exceeds the reach of direct connections by a large factor (Figure 3*D*). As the network approaches instability, which corresponds to the spectral bound *R* going to one, the effective decay constant diverges (Figure 3*D inset*) and so does the range of covariances.

Our population-resolved theoretical analysis, furthermore, shows that the larger the spectral bound the more similar the decay constants between different populations, with only marginal differences for *R* ≲ 1 (Figure 3*E*). This holds strictly if connection weights only depend on the type of the presynaptic neuron but not on the type of the postsynaptic neuron. Moreover, we find a relation between the squared effective decay constants and the squared anatomical decay constants of the form

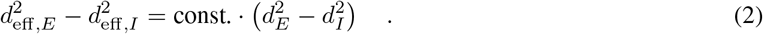

This relation is independent of the eigenvalues of the effective connectivity matrix, as the constant of order 𝒪 (1) does only depend on the choice of the connectivity profile. For *R* 1, this means that even though the absolute value of both effective length scales on the left hand side is large, their relative difference is small because it equals the small difference of anatomical length scales on the right hand side.

### Pairwise covariances in motor cortex decay on a millimeter scale

To check if these predictions are confirmed by the data from macaque motor cortex, we first observe that, indeed, covariances in the resting state show a large dispersion over almost all distances on the Utah array (Figure 4). Moreover, the variance of covariances agrees well with the predicted exponential law: Performing an exponential fit reveals length constants above one millimeter. These large length constants have to be compared to the spatial reach of direct connections, which is about an order of magnitude shorter, in the range of 100 − 400 *µ*m (*22*), so below the 400 *µ*m inter-electrode distance of the Utah array. The shallow decay of the variance of covariances is, next to the broad distribution of covariances, a second indication that the network is in the dynamically balanced critical regime, in line with the prediction by Equation (1).

**Figure 4:**
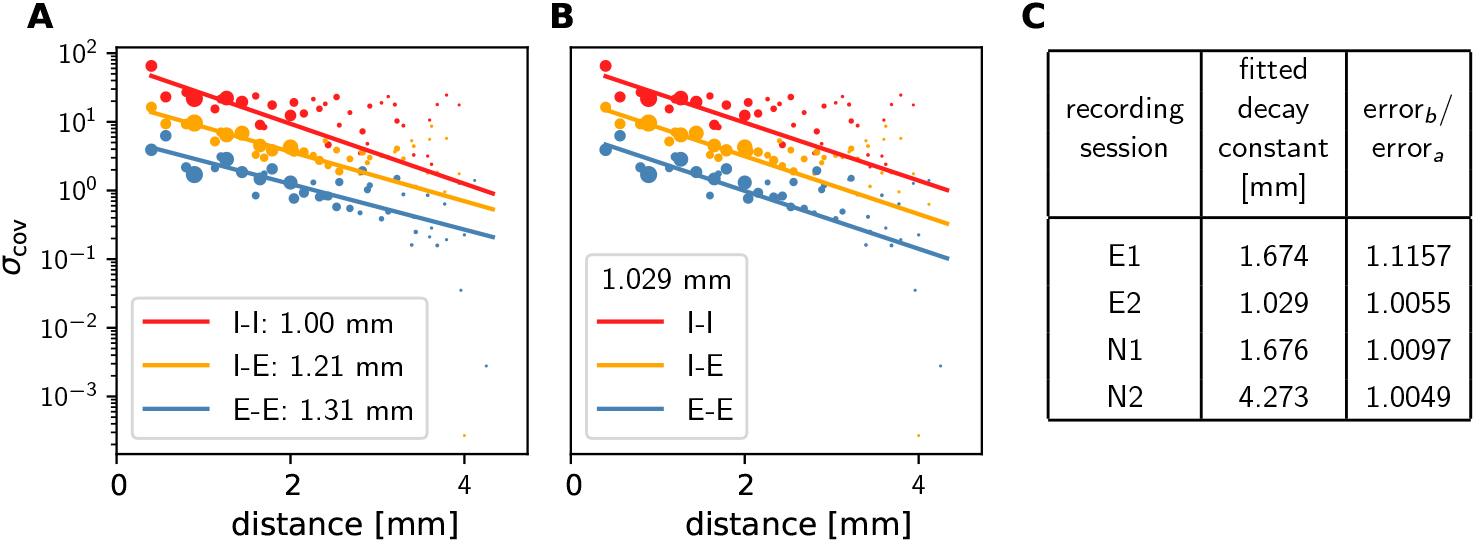
Long-range covariances in macaque motor cortex. Variance of covariances as a function of distance. (*A*) Population-specific exponential fits (lines) to variances of covariances (dots) in session E2, with fitted decay constants indicated in the legend (I-I: putative inhibitory neuron pairs, I-E: inhibitory-excitatory, E-E: excitatory pairs). Dots show the empirical estimate of the variance of the covariance distribution for each distance. Size of the dots represents relative count of pairs per distance and was used as weighting factor for the fits to compensate for uncertainty at large distances, where variance estimates are based on fewer samples. Mean squared error 2.918. (*B*) Population-specific exponential fits (lines) analogous to (*A*), with slopes constrained to be identical. This procedure yields a single fitted decay constant indicated in the legend. Mean squared error 2.934. (*C*) Table listing decay constants fitted as in (*B*) for all recording sessions and the ratios between mean squared errors of the fits obtained in procedures *B* and *A*. The source data and code used to create this figure are provided as fig4.zip.

The population-resolved fits to the data show a larger length constant for excitatory covariances than for inhibitory ones (Figure 4*A*). This is qualitatively in line with the prediction of Equation (2) given the typically longer reach of excitatory connections compared to inhibitory ones (*36, 37*). In the dynamically balanced critical regime, however, the predicted difference in slope for all three fits is practically negligible. Therefore, we performed a second fit where the slope of the three exponentials is constrained to be identical (Figure 4*B*). The error of this fit is only marginally larger than the ones of fitting individual slopes (Figure 4*C*). This shows that differences in slopes are hardly detectable given the empirical evidence, thus confirming the predictions of the theory given by Equation (1) and Equation (2).

### Firing rates alter connectivity-dependent covariance patterns

Since covariances measure the coordination of temporal fluctuations around the individual neurons’ mean firing rates, they are determined by how strong a neuron transmits such fluctuations from input to output (*38*). To leading order this is explained by linear response theory (*2, 35, 24, 23*): How strongly a neuron reacts to a small change in its input depends on its dynamical state, foremost the mean and variance of its total input, called “working point” in the following. If a neuron receives almost no input, a small perturbation in the input will not be able to make the neuron fire. If the neuron receives a large input, a small perturbation will not change the firing rate either, as the neuron is already saturated. Only in the intermediate regime the neuron is susceptible to small deviations of the input. Mathematically, this behavior is described by the gain of the neuron, which is the derivative of the input-output relation (*38*). Due to the non-linearity of the input-output relation, the gain is vanishing for very small and very large inputs and non-zero in the intermediate regime. How strongly a perturbation in the input to one neuron affects one of the subsequent neurons therefore not only depends on the synaptic weight ***J*** but also on the gain ***S*** and thereby the working point. This relation is captured by the effective connectivity ***W*** = ***S* · *J***. What is the consequence of the dynamical interaction among neurons depending on the working point? Can it be used to reshape the low-dimensional manifold, the collective coordination between neurons? The first part of this study finds that long-range coordination can be achieved in a network with short-range random connections if effective connections are sufficiently strong. Alteration of the working point, for example by a different external input, can affect the covariance structure: The pattern of coordination between individual neurons can change, even though the anatomical connectivity remains the same. In this way, routing of information through the network can be adapted dynamically on a mesoscopic scale. This is a crucial difference of such coordination as opposed to coordination imprinted by complex but static connection patterns.

We first illustrate this concept by simulations of a network of nonlinear rate model neurons. For independent and stationary external inputs covariances between neurons are solely generated inside the network via the recurrent connectivity. External inputs only have an indirect impact on the covariance structure by setting the working point of the neurons.

We create two networks with identical structural connectivity and identical external input fluctuations (Figure 5*A*). Small differences in mean external inputs between corresponding neurons in the two networks create slightly different gains and firing rates and thereby differences in effective connectivity and covariances. Since mean external inputs are drawn from the same distribution in both networks (Figure 5*B*), the overall distributions of firing rates and covariances across all neurons are very similar (Figure 5*E1,F2*). But individual neurons’ firing rates do differ (Figure 5*E2*). The resulting change of the neurons’ working points substantially affects the covariance patterns (Figure 5*F2*): Differences in firing rates and covariances between the two networks are significantly larger than the differences within the same network across two different time periods (Figure 5*C*). The larger the spectral bound, the more sensitive are the intrinsically generated covariances to the changes in firing rates (Figure 5*D*). Thus, a small offset of individual firing rates is an effective parameter to control network-wide coordination among neurons. As the input to the local network can be changed momentarily, we predict that in the dynamically balanced critical regime coordination patterns should be highly dynamic.

**Figure 5:**
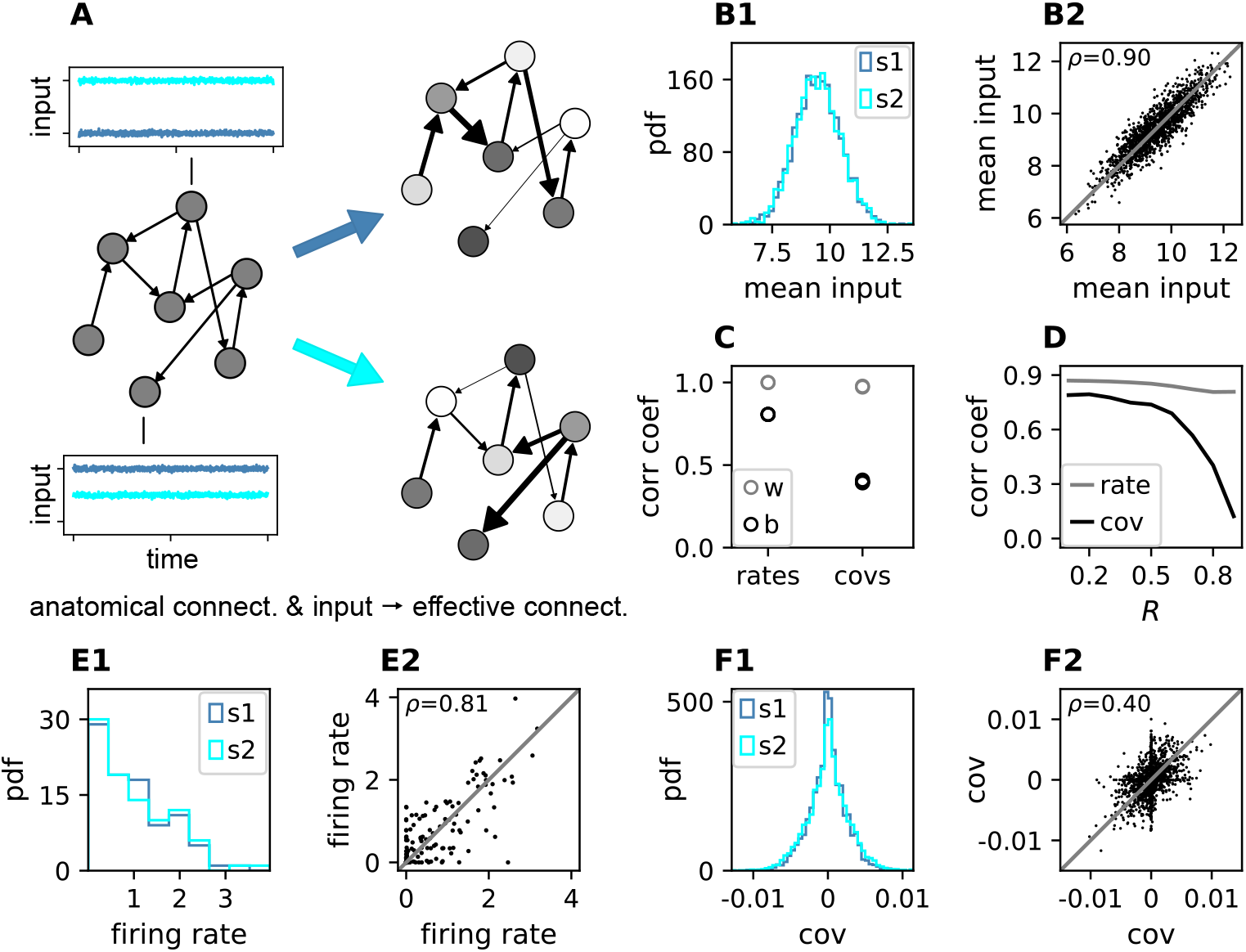
Changes in effective connectivity modify coordination patterns. (*A*) Two sparse random networks with identical structural connectivity (left network sketch) but different inputs (depicted by insets), which results in different firing rates (grayscale in right network sketches) and therefore different effective connectivities (thickness of connections). Parameters can be found in table 3 in Appendix 1. (*B1*) Histogram of input currents across neurons for the two networks (s1 and s2). (*B2*) Scatter plot of inputs to subset of 1500 corresponding neurons in the first and the second network (Pearson correlation coefficient *ρ* = 0.90). (*C*) Correlation coefficients of rates and of covariances between the networks (b, black) and within the same network (w, gray; simulation was split into two halves). (*D*) Correlation coefficient of rates (gray) and covariances (black) between the two networks as a function of the spectral bound *R*. (*E1*) Distribution of rates in the two networks (excluding silent neurons with |rate| < 10^−3^). (*E2*) Scatter plot of rates in the first compared to the second network (Pearson correlation coefficient *ρ* = 0.81). (*F1*) Distribution of covariances in the two networks (excluding silent neurons). (*F2*) Scatter plot of sample of 5000 covariances in first compared to the second network (Pearson correlation coefficient *ρ* = 0.40). Other parameters: number of neurons *N* = 2000, connection probability *p* = 0.1, spectral bound for panels *B, C, E, F* is *R* = 0.8. The source data and code used to create this figure are provided as fig5.zip.

### Coordination patterns in motor cortex depend on behavioral context

In order to test the model prediction in experimental data, we analyze massively-parallel spiking activity from macaque motor cortex, recorded during a reach-to-grasp experiment (*39, 40*). In contrast to the resting state, where the animal was in an idling state, here the animal is involved in a complex task with periods of different cognitive and behavioral conditions (Figure 6*A*). We compare two epochs in which the animal is requested to wait and is sitting still but which differ in cognitive conditions. The first epoch is a starting period (S), where the monkey has self-initiated the behavioral trial and is attentive because it is expecting a cue. The second epoch is a preparatory period (P), where the animal has just received partial information about the upcoming trial and is waiting for the missing information and the GO signal to initiate the movement. In both epochs the neuronal firing rates are stationary, likely due to the absence of arm movement (see Appendix 1 section 2). The overall distributions of the firing rates in the different epochs are comparable (Figure 6*C*) but are distributed differently across the individual neurons. By comparing the firing rates across neurons between disjoint sub-periods of any epoch (e.g. S1-S2 or S1-P2), we observe that the firing rate compositions across neurons change more strongly across the two epochs (Sx-Py) than within each of them (Figure 6*E*). This holds for data from five/eight different recording sessions from different recording days for monkey E/N. Similarly, the covariance values change, though here the changes are even more pronounced (Figure 6*F*). This is in line with our prediction for a network whose effective connectivity has a large spectral bound, in the critically balanced state. In particular, the theory predicts different coordination patterns between neurons on the mesoscopic scale (range of a Utah array), which is indeed observed in the two states S and P (Figure 6*B*). The coordination between neurons is thus considerably reshaped by the behavioral condition.

**Figure 6:**
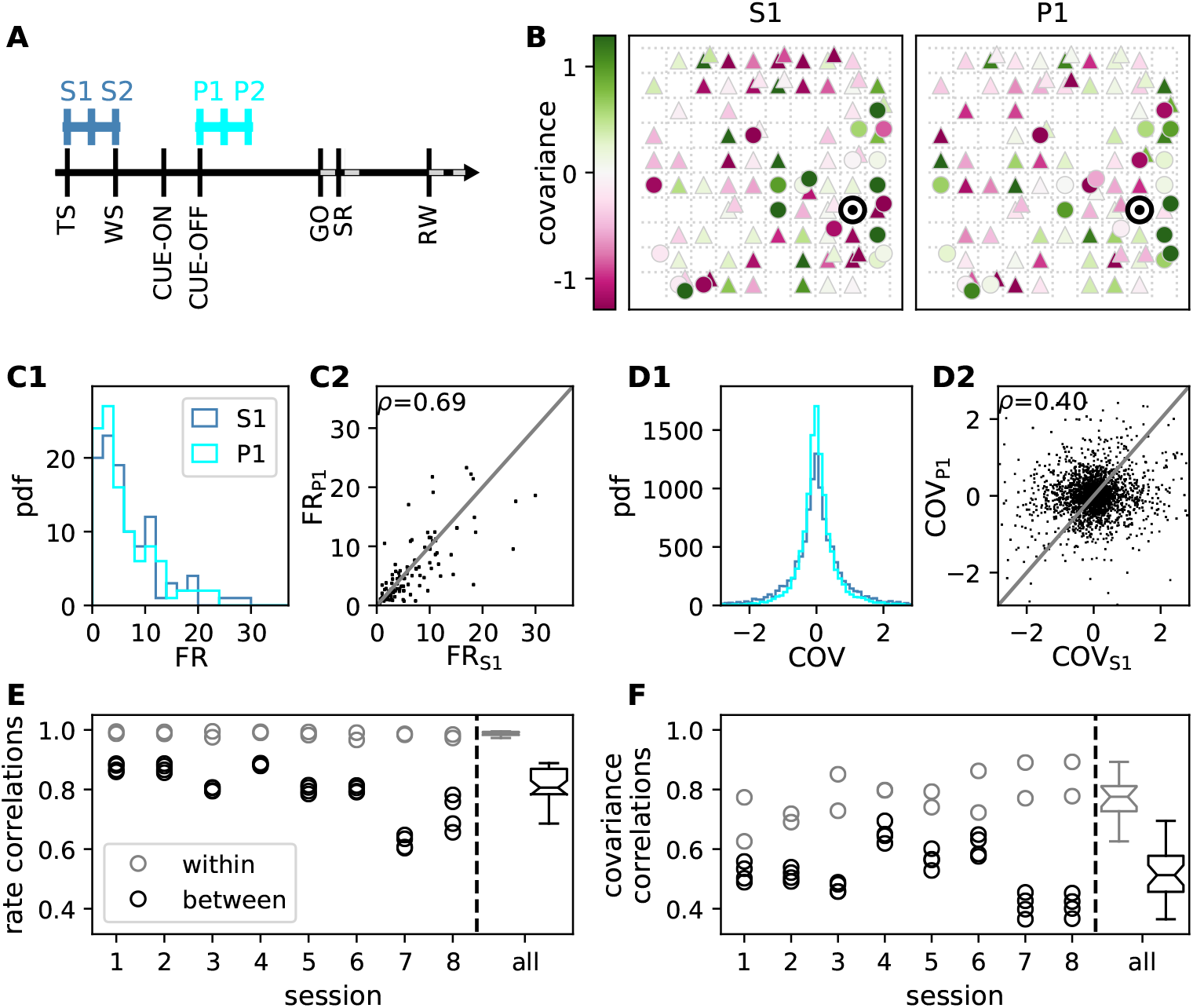
Behavioral condition reshapes mesoscopic neuronal coordination. (*A*) Trial structure of the reach-to-grasp experiment (*39*). Blue segments above the time axis indicate data pieces at trial start (dark blue: S1, S2) and during the preparatory period (light blue: P1, P2). (*B*) Salt-and-pepper structure of covariance during two different epochs (S1 and P1) of one recording session of monkey N (151 trials, 106 single units, cf. Figure HYPERLINK \l “bookmark0” 1 for recording setup). For some neurons the covariance completely reverses, while in the others it does not change. Inhibitory reference neuron indicated by black circle. (*C1*) Distributions of firing rates during S1 and P1. (*C2*) Scatter plot comparing firing rates in S1 and P1 (Pearson correlation coefficient *ρ* = 0.69). (*D1*/*D2*) Same as panels *C1*/*C2*, but for covariances (Pearson correlation coefficient *ρ* = 0.40). (*E*) Correlation coefficient of firing rates across neurons in different epochs of a trial for eight recorded sessions. Correlations between sub-periods of the same epoch (S1-S2, P1-P2; within-epoch, gray) and between sub-periods of different epochs (Sx-Py; between-epochs, black). Box plots to the right of the black dashed line show distributions obtained after pooling across all analyzed recording sessions per monkey. The line in the center of each box represents the median, box’s area represents the interquartile range, and the whiskers indicate minimum and maximum of the distribution (outliers excluded). Those distributions differ significantly (Student t-test, two-sided, *p* ≪ 0.001). (*F*) Correlation coefficient of covariances, analogous to panel e. The distributions of values pooled across sessions also differ significantly (Student t-test, two-sided, *p* ≪ 0.001). For details of the statistical tests, see Methods. Details on number of trials and units in each recording session are provided in Appendix 1-table 2. The source data and code used to create this figure are provided as fig6.zip.

## Discussion

In this study, we investigate coordination patterns of many neurons across mesoscopic distances in macaque motor cortex. We show that these patterns have a salt-and-pepper structure, which can be explained by a network model with a spatially dependent random connectivity operating in a dynamically balanced critical state. In this state, cross-covariances are shaped by a large number of parallel, multi-synaptic pathways, leading to interactions reaching far beyond the range of direct connections. Strikingly, this coordination on the millimeter scale is only visible if covariances are resolved on the level of individual neurons; the population mean of covariances quickly decays with distance and is overall very small. In contrast, the variance of covariances is large and predominantly decreases exponentially on length scales of up to several millimeters, even though direct connections typically only reach a few hundred micrometers.

Since the observed coordination patterns are determined by the effective connectivity of the network, they are dynamically controllable by the network state; for example, due to modulations of neuronal firing rates. Massively parallel recordings in macaque motor cortex during resting state and in different epochs of a reach-to-grasp task confirm this prediction. Simulations indeed exhibit a high sensitivity of coordination patterns to weak modulations of the individual neurons’ firing rates, providing a plausible mechanism for these dynamic changes.

Models of balanced networks have been investigated before (*26*,41,3,23) and experimental evidence for cortical networks operating in the balanced state is overwhelming (*42, 43, 44*). Excess of inhibition in such networks yields stable and balanced population-averaged activities as well as low average covariances (*23*). Recently the notion of balance has been combined with criticality in the dynamically balanced critical state that results from large heterogeneity in the network connectivity (*15*). Here we focus on another ubiquitous property of cortical networks, their spatial organization, and study the interplay between balance, criticality, and spatial connectivity in networks of excitatory and inhibitory neurons. We show that in such networks, heterogeneity generates disperse covariance structures between individual neurons on large length-scales with a salt-and-pepper structure.

Spatially organized balanced network models have been investigated before in the limit of infinite network size, as well as under strong and potentially correlated external drive (*5, 30*). In this limit, intrinsically generated covariances can be neglected and population-averaged covariances fulfill a linear equation, called the “balance condition”, that predicts a non-monotonous change of population-averaged covariances with distance (*45*). In contrast, we here consider covariances on the level of individual cells in finite-size networks with weak inputs or in resting state conditions. In such a scenario, covariances have been shown to be predominantly generated locally rather than from external inputs (*4, 15*). Analyses on the single-neuron level go beyond the balance condition and require the use of field-theoretical techniques to capture the heterogeneity in the network (*15*).

The analysis performed here requires a theory for disordered systems with spatially non-homogeneous coupling to analyze excitatory-inhibitory random networks on a two-dimensional sheet with spatially decaying connection probabilities. This new theory allows us to derive expressions for the spatial decay of the variance of covariances. We primarily evaluate these expressions in the long-range limit, which agrees well with simulations for distances *x* > 2*d* ∼ 𝒪 (1 mm), which is fulfilled for most distances on the Utah array (cf. Figure 3, Appendix 1-figure 4). For these distances we find that the decay of covariances is dominated by a simple exponential law. Unexpectedly, its decay constant is essentially determined by only two measures, the spectral bound of the effective connectivity, and the length scale of direct connections. The length scale of covariances diverges when approaching the breakdown of linear stability. In this regime, differences in covariances induced by differences in length scales of excitatory and inhibitory connections become negligible. The predicted emergence of a single length scale of covariances is consistent with our data.

This study focuses on local and isotropic connection profiles to show that long-range coordination does not rely on specific connection patterns but can result from the network state alone. Alternative explanations for long-range coordination are based on specifically imprinted network structures: Anisotropic local connection profiles have been studied and shown to create spatio-temporal sequences (*46*). Likewise, embedded excitatory feed-forward motifs and cell assemblies via long-range patchy connections can create positive covariances at long distances (*47, 48*). These structures are, however, static: A change in gain of the neurons will either strengthen or weaken the specific activity propagation, but it will not lead to new pathways of propagation within the network and therefore not cause significantly different coordination patterns that we see in our data. The static impact of these connectivity structures on covariances could in principle be included in the presented formalism. Long-range coordination can also be created from short-range connections with random orientations of anisotropic local connection profiles (*14*). This finding can be linked to the emergence of tuning maps in the visual cortex. The mechanism is similar to ours in that it uses nearly linearly unstable modes that are determined by spatial connectivity structures and heterogeneity. Given the different source of heterogeneity, the modes and corresponding covariance patterns are different from the ones discussed here: Starting from fully symmetric networks with corresponding symmetric covariance patterns, (*14*) found that increasing heterogeneity yields more randomized, but still patchy regions of positive and negative covariances that are in line with low-dimensional activity patterns found in visual cortex. In motor cortex we instead find salt-and-pepper patterns that can be explained in terms of heterogeneity through sparsity. We provide the theoretical basis and explicit link between connectivity eigenspectra and covariances and show that heterogeneity through sparsity is sufficient to generate the dynamically balanced critical state as a simple explanation for the broad distribution of covariances in motor cortex, the salt-and-pepper structure of coordination, its long spatial range, and its sensitive dependence on the network state.

What are possible functional implications of the coordination on mesoscopic scales? Recent work demonstrated activity in motor cortex to be organized in low-dimensional manifolds (*6, 7, 8*). Dimensionality reduction techniques, such as PCA or GPFA (*49*), employ covariances to expose a dynamical repertoire of motor cortex that is comprised of neuronal modes. Previous work started to analyze the relation between the dimensionality of activity and connectivity (*16, 17, 18, 15, 20, 21*), but only in spatially unstructured networks, where each neuron can potentially be connected to any other neuron. The majority of connections within cortical areas, however, stems from local axonal arborizations (*22*). Here we add this biological constraint and demonstrate that these networks, too, support a dynamically balanced critical state. This state in particular exhibits neural modes which are spanned by neurons spread across the experimentally observed large distances. In this state a small subset of modes that are close to the point of instability dominates the variability of the network activity and thus spans a low-dimensional neuronal manifold. As opposed to specifically designed connectivity spectra via plasticity mechanisms (*50*) or low-rank structures embedded into the connectivity (*18*), the dynamically balanced critical state is a mechanism that only relies on the heterogeneity which is inherent to sparse connectivity and abundant across all brain areas.

While we here focus on covariance patterns in stationary activity periods, the majority of recent works studied transient activity during motor behavior (*6*). How are stationary and transient activities related? During stationary ongoing activity states, covariances are predominantly generated intrinsically (*4*). Changes in covariance patterns therefore arise from changes in the effective connectivity via changes in neuronal gains, as demonstrated here in the two periods of the reach-to-grasp experiment and in our simulations for networks close to criticality (Figure 5*D*). During transient activity, on top of gain changes, correlated external inputs may directly drive specific neural modes to create different motor outputs, thereby restricting the dynamics to certain subspaces of the manifold. In fact, (*51*) reported that the covariance structures during movement preparation and movement execution are unrelated and corresponding to orthogonal spaces within a larger manifold. Also (*52*) studied auditory and somatosensory cortices of awake and anesthetized rats during spontaneous and stimulus-evoked conditions and found that neural modes of stimulus-evoked activity lie in subspaces of the neural manifold spanned by the spontaneous activity. Similarly, visual areas V1 and V2 seem to exploit distinct subspaces for processing and communication (*53*), and motor cortex uses orthogonal subspaces capturing communication with somatosensory cortex or behavior-generating dynamics (*54*). (*7*) further showed that manifolds are not identical, but to a large extent preserved across different motor tasks due to a number of task-independent modes. This leads to the hypothesis that the here described mechanism for long-range cooperation in the dynamically balanced critical state provides the basis for low-dimensional activity by creating such spatially extended neural modes, whereas transient correlated inputs lead to their differential activation for the respective target outputs. The spatial spread of the neural modes thereby leads to a distributed representation of information that may be beneficial to integrate information into different computations that take place in parallel at various locations. Further investigation of these hypotheses is an exciting endeavor for the years to come.

## Methods

### Experimental Design and Statistical Analysis

Two adult macaque monkeys (monkey E - female, and monkey N - male) are recorded in behavioral experiments of two types: resting state and reach-to-grasp. The recordings of neuronal activity in motor and pre-motor cortex (hand/arm region) are performed with a chronically implanted 4×4 mm^2^ Utah array (Blackrock Microsystems). Details on surgery, recordings, spike sorting and classification of behavioral states can be found in (*40, 55, 39, 56*). All animal procedures were approved by the local ethical committee (C2EA 71; authorization A1/10/12) and conformed to the European and French government regulations.

#### Resting state data

During the resting state experiment, the monkey is seated in a primate chair without any task or stimulation. Registration of electrophysiological activity is synchronized with a video recording of the monkey’s behavior. Based on this, periods of “true resting state” (RS), defined as no movements and eyes open, are chosen for the analysis. Eye movements and minor head movements are included. Each monkey is recorded twice, with a session lasting approximately 15 and 20 min for monkeys E (sessions E1 and E2) and N (sessions N1 and N2), respectively, and the behavior is classified by visual inspection with single second precision, resulting in 643 and 652 s of RS data for monkey E and 493 and 502 s of RS data for monkey N.

#### Reach-to-grasp data

In the reach-to-grasp experiment, the monkeys are trained to perform an instructed delayed reach-to-grasp task to obtain a reward. Trials are initiated by a monkey closing a switch (TS, trial start). After 400 ms a diode is illuminated (WS, warning signal), followed by a cue after another 400 ms(CUE-ON), which provides partial information about the upcoming trial. The cue lasts 300 ms and its removal (CUE-OFF) initiates a 1 s preparatory period, followed by a second cue, which also serves as GO signal. Two epochs, divided into 200 ms sub-periods, within such defined trials are chosen for analysis: the first 400 ms after TS (starting period, S1 and S2), and the 400 ms directly following CUE-OFF (preparatory period, P1 and P2) (cf. Figure 6a). Five selected sessions for monkey E and eight for monkey N provide a total of 510 and 1111 correct trials, respectively. For detailed numbers of trials and single units per recording session see Appendix 1-table 2.

#### Separation of putative excitatory and inhibitory neurons

Offline spike-sorted single units (SUs) are separated into putative excitatory (broad-spiking) and putative inhibitory (narrow-spiking) based on their spike waveform width (*57, 58, 59, 60, 61*). The width is defined as the time (number of data samples) between the trough and peak of the waveform. Widths of all average waveforms from all selected sessions (both resting state and reach-to-grasp) per monkey are collected. Thresholds for “broadness” and “narrowness” are chosen based on the monkey-specific distribution of widths, such that intermediate values stay unclassified. For monkey E the thresholds are 0.33 ms and 0.34 ms, and for monkey N 0.40 ms and 0.41 ms. Next, a two step classification is performed session by session. Firstly, the thresholds are applied to average SU waveforms. Secondly, the thresholds are applied to SU single waveforms and a percentage of single waveforms pre-classified as the same type as the average waveform is calculated. SU for which this percentage is high enough are marked classified. All remaining SUs are grouped as unclassified. We verify the robustness of our results with respect to changes in the spike sorting procedure in Appendix 1 section 1.

Synchrofacts, i.e., spike-like synchronous events across multiple electrodes at the sampling resolution of the recording system (1*/*30 ms) (*62*), are removed. In addition, only SUs with a signal-to-noise ratio (*63*) of at least

2.5 and a minimal average firing rate of 1 Hz are considered for the analysis, to ensure enough and clean data for valid statistics.

#### Statistical analysis

All RS periods per resting state recording are concatenated and binned into 1 s bins. Next, pairwise covariances of all pairs of SUs are calculated according to the following formula:

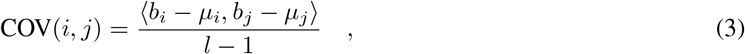

with *b*_*i*_, *b*_*j*_ - binned spike trains, *µ*_*i*_, *µ*_*j*_ being their mean values, *l* the number of bins, and *x, y* the scalar product of vectors *x* and *y*. Obtained values are broadly distributed, but low on average in every recorded session: in session E1 E-E pairs: 0.19 ± 1.10 (M ± SD), E-I: 0.24 ± 2.31, I-I: 0.90 ± 4.19, in session E2 E-E: 0.060 ± 1.332, E-I 0.30 ± 2.35, I-I 1.0 ± 4.5, in session N1 E-E 0.24 ± 1.13, E-I 0.66 ± 2.26, I-I 2.4± 4.9, in session N2 E-E 0.41 ± 1.47, E-I 1.0 ± 3.1, I-I 3.9 ± 7.3.

To explore the dependence of covariance on the distance between the considered neurons, the obtained values are grouped according to distances between electrodes on which the neurons are recorded. For each distance the average and variance of the obtained distribution of cross-covariances is calculated. The variance is additionally corrected for bias due to a finite number of measurements (*15*). In most of cases, the correction does not exceed 0.01%.

In the following step, exponential functions 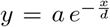 are fitted to the obtained distance-resolved variances of cross-covariances (*y* corresponding to the variance and *x* to distance between neurons), which yields a pair of values (*a, d*). The least squares method implemented in the Python scipy.optimize module (SciPy v.1.4.1) is used. Firstly, three independent fits are performed to the data for excitatory-excitatory, excitatory-inhibitory, and inhibitory-inhibitory pairs. Secondly, analogous fits are performed, with the constraint that the decay constant *d* should be the same for all three curves.

Covariances in the reach-to-grasp data are calculated analogously but with different time resolution. For each chosen sub-period of a trial, data are concatenated and binned into 200 ms bins, meaning that the number of spikes in a single bin corresponds to a single trial. The mean of these counts normalized to the bin width gives the average firing rate per SU and sub-period. The pairwise covariances are calculated according to Equation (3). To assess the similarity of neuronal activity in different periods of a trial, Pearson product-moment correlation coefficients are calculated on vectors of SU-resolved rates and pair-resolved covariances. Correlation coefficients from all recording sessions per monkey are separated into two groups: using sub-periods of the same epoch (*within*-epoch), and using sub-periods of different epochs of a trial (*between*-epochs). These groups are tested for differences with significance level *α* = 0.05. Firstly, to check if the assumptions for parametric tests are met, the normality of each obtained distribution is assessed with a Shapiro-Wilk test, and the equality of variances with an F-test. Secondly, a t-test is applied to compare within- and between-epochs correlations of rates or covariances. Since there are two *within* and four *between* correlation values per recording session, the number of degrees of freedom equals: *df* = (*N*_sessions_ · 2 − 1) + (*N*_sessions_ 4 − 1), which is 28 for monkey E and 46 for monkey N. To estimate the confidence intervals for obtained differences, the mean difference between groups *m* and their pooled standard deviation *s* are calculated for each comparison the number of *within-* and *between*-epoch correlation coefficient values, respectively.

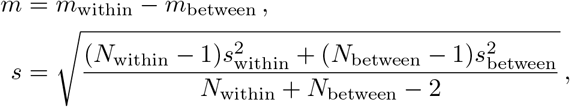

with *m*_*within*_ and *m*_*between*_ being the mean, *s*_*within*_ and *s*_*between*_ the standard deviation and *N*_*within*_ and *N*_*between*_ the number of within- and between-epoch correlation coefficient values, respectively.

This results in 95% confidence intervals *m* ± *t*(*df*) · *s* of 0.192 ± 0.093 for rates and 0.32 ± 0.14 for covariances in monkey E and 0.19 ± 0.14 for rates and 0.26 ± 0.17 for covariances in monkey N.

For both monkeys the *within*-epoch rate-correlations distribution does not fulfill the normality assumption of the t-test. We therefore perform an additional non-parametric Kolmogorov-Smirnov test for the rate comparison. The differences are again significant; for monkey E *D* = 1.00, *p* = 6.66 · 10^−8^; for monkey N *D* = 1.00, *p* = 8.87 · 10^−13^.

For all tests we use the implementations from the Python scipy.stats module (SciPy v.1.4.1).

### Mean and Variance of Covariances for a Two-Dimensional Network Model with Excitatory and Inhibitory Populations

The mean and variance of covariances are calculated for a two-dimensional network consisting of one excitatory and one inhibitory population of neurons. The connectivity profile *p*(***x***), describing the probability of a neuron having a connection to another neuron at distance ***x***, decays with distance. We assume periodic boundary conditions and place the neurons on a regular grid (Figure HYPERLINK \l “bookmark3” 3*A*), which imposes translation and permutation symmetries that enable the derivation of closed-form solutions for the distance-dependent mean and variance of the covariance distribution. These simplifying assumptions are common practice and simulations show that they do not alter the results qualitatively.

Our aim is to find an expression for the mean and variance of covariances as functions of distance between two neurons. While the theory in (*15*) is restricted to homogeneous connections, understanding the spatial structure of covariances here requires us to take into account the spatial structure of connecti<u>vity</u>. Field-theoretic methods allow us to obtain expressions for the mean covariance 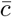 and variance of covariance 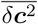

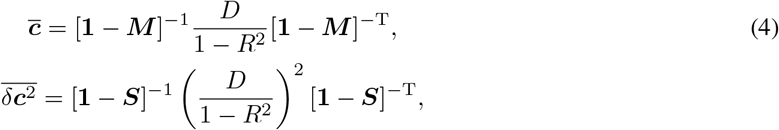

with identity matrix **1**, mean ***M*** and variance ***S*** of connectivity matrix ***W***, input noise strength *D*, and spectral bound *R*. Since ***M*** and ***S*** have a similar structure, the mean and variance can be derived in the same way, which is why we only consider variances in the following.

To simplify Equation (4), we need to find a basis in which ***S***, and therefore also ***A*** = **1** − ***S***, is diagonal. Due to invariance under translation, the translation operators ***T*** and the matrix ***S*** have common eigenvectors, which can be derived using that translation operators satisfy ***T*** ^*N*^ = 1, where *N* is the number of lattice sites in *x*- or *y*-direction (see Appendix 1). Projecting onto a basis of these eigenvectors shows that the eigenvalues *s*_***k***_ of ***S*** are given by a discrete two-dimensional Fourier transform of the connectivity profile

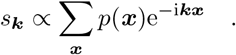

Expressing ***A***^−1^ in the eigenvector basis yields ***A***^−1^(***x***) = **1** + ***B***(***x***), where ***B***(***x***) is a discrete inverse Fourier transform of the kernel *s*_***k***_*/*(1 − *s*_***k***_). Assuming a large network with respect to the connectivity profiles allows us to take the continuum limit

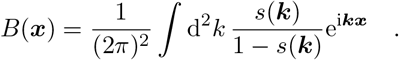

As we are only interested in the long-range behavior, which corresponds to |***x***| → ∞, or |***k***| →0, respectively, we can approximate the Fourier kernel around |***k***| ≈ 0 by a rational function, quadratic in the denominator, using a Padé approximation. This allows us to calculate the integral which yields

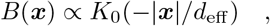

where *K*_0_(*x*) denotes the modified Bessel function of second kind and zeroth order (*64*), and the effective decay constant *d*_eff_ is given by Equation (1). In the long-range limit the modified Bessel function behaves like

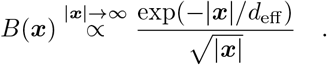

Writing Equation (4) in terms of *B*(***x***) gives

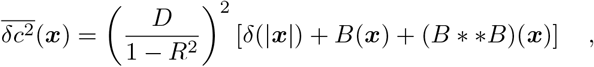

with the double asterisk denoting a two-dimensional convolution. (*B* * * *B*)(***x***) is a function proportional to the modified Bessel function of second kind and first order (*64*), which has the long-range limit

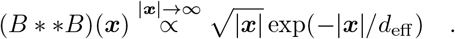

Hence, the effective decay constant of the variances is given by *d*_eff_. Note that further details of the above derivation can be found in the Appendix 1 section 3-11.

## Code and Data availability

All code and data required to reproduce the figures are available in a public zenodo repository at http://doi.org/10.5281/zenodo.4686549.

## Acknowledgments

This work was partially supported by HGF young investigator’s group VH-NG-1028, European Union’s Horizon 2020 research and innovation program under Grant agreements No. 785907 (Human Brain Project SGA2) and No. 945539 (Human Brain Project SGA3), ANR grant GRASP and partially funded by the Deutsche Forschungsgemeinschaft (DFG, German Research Foundation) - 368482240/GRK2416. We are grateful to our colleagues in the NEST and Elephant developer communities and for continuous collaboration. All network simulations were carried out with NEST 2.20.0 (http://www.nest-simulator.org). All data analyses performed with Elephant (https://neuralensemble.org/elephant/). We thank Sebastian Lehmann for help with the design of the figures.

## References

1. A. P. Georgopoulos, R. Caminiti, J. F. Kalaska, J. T. Massey, Experimental Brain Research [Suppl.] 7, 327 (1983).

2. I. Ginzburg, H. Sompolinsky, Phys. Rev. E 50, 3171 (1994).

3. A. Renart, et al., Science 327, 587 (2010).

4. M. Helias, T. Tetzlaff, M. Diesmann, PLOS Comput. Biol. 10, e1003428 (2014).

5. R. Rosenbaum, B. Doiron, Physical Review X 4, 021039 (2014).

6. J. A. Gallego, M. G. Perich, L. E. Miller, S. A. Solla, Neuron 94, 978 (2017).

7. J. A. Gallego, et al., Nature Communications 9, 1 (2018).

8. J. A. Gallego, M. G. Perich, R. H. Chowdhury, S. A. Solla, L. E. Miller, Nat. Neurosci. 23, 260 (2020).

9. D. Sussillo, M. M. Churchland, M. T. Kaufman, K. V. Shenoy, Nat. Neurosci. 18, 1025 (2015).

10. L. F. Abbott, K. Rajan, H. Sompolinsky, The Dynamic Brain: An Exploration of Neuronal Variability and its Functional Significance (2011), pp. 65–82.

11. L. Mazzucato, A. Fontanini, G. La Camera, Frontiers in systems neuroscience 10, 11 (2016).

12. P. Gao, et al., bioRxiv p. 214262 (2017).

13. C. Stringer, M. Pachitariu, N. Steinmetz, M. Carandini, K. D. Harris, Nature 571, 361 (2019).

14. G. B. Smith, B. Hein, D. E. Whitney, D. Fitzpatrick, M. Kaschube, Nat. Neurosci. 21, 1600 (2018).

15. D. Dahmen, S. Grün, M. Diesmann, M. Helias, Proc. Nat. Acad. Sci. USA 116, 13051 (2019).

16. J. Aljadeff, M. Stern, T. Sharpee, Phys. Rev. Lett. 114, 088101 (2015).

17. J. Aljadeff, D. Renfrew, M. Vegué, T. O. Sharpee, Phys. Rev. E 93, 022302 (2016).

18. F. Mastrogiuseppe, S. Ostojic, Neuron 99, 609 (2018).

19. S. Recanatesi, G. K. Ocker, M. A. Buice, E. Shea-Brown, PLOS Comput. Biol. 15, 1 (2019).

20. D. Dahmen, et al., bioRxiv (2020).

21. Y. Hu, H. Sompolinsky, bioRxiv (2020).

22. P. Schnepel, A. Kumar, M. Zohar, A. Aertsen, C. Boucsein, Cerebral Cortex 25, 3818 (2015).

23. T. Tetzlaff, M. Helias, G. T. Einevoll, M. Diesmann, PLOS Comput. Biol. 8, e1002596 (2012).

24. V. Pernice, B. Staude, S. Cardanobile, S. Rotter, PLOS Comput. Biol. 7, e1002059 (2011).

25. V. Pernice, B. Staude, S. Cardanobile, S. Rotter, Phys. Rev. E 85, 031916 (2012).

26. C. van Vreeswijk, H. Sompolinsky, Science 274, 1724 (1996).

27. D. J. Amit, N. Brunel, Cereb. Cortex 7, 237 (1997).

28. C. van Vreeswijk, H. Sompolinsky, Neural Comput. 10, 1321 (1998).

29. R. Darshan, C. van Vreeswijk, D. Hansel 8, 031072 (2018).

30. C. Baker, C. Ebsch, I. Lampl, R. Rosenbaum, Phys. Rev. E 99, 052414 (2019).

31. J. Trousdale, Y. Hu, E. Shea-Brown, K. Josic, PLOS Comput. Biol. 8, e1002408 (2012).

32. B. A. W. Brinkman, F. Rieke, E. Shea-Brown, M. A. Buice, PLOS Comput. Biol. 14, e1006490 (2018).

33. R. Kobayashi, et al., Nature Communications 10, 1 (2019).

34. J. De la Rocha, B. Doiron, E. Shea-Brown, J. Kresimir, A. Reyes, Nature 448, 802 (2007).

35. B. Lindner, B. Doiron, A. Longtin, Phys. Rev. E 72, 061919 (2005).

36. M. W. Reimann, A.-L. Horlemann, S. Ramaswamy, E. B. Muller, H. Markram, Cereb. Cortex 27, 4570 (2017).

37. Y. Hao, A. Riehle, T. G. Brochier, Frontiers in neural circuits 10, 104 (2016).

38. M. Abeles, Corticonics: Neural Circuits of the Cerebral Cortex (Cambridge University Press, Cambridge, 1991), first edn.

39. T. Brochier, et al., Scientific Data 5, 180055 (2018).

40. A. Riehle, S. Wirtssohn, S. Grün, T. Brochier, Frontiers in Neural Circuits 7, 48 (2013).

41. N. Brunel, Journal of Computational Neuroscience 8, 183 (2000).

42. M. Okun, I. Lampl, Nat. Neurosci. 11, 535 (2008).

43. K. Reinhold, A. D. Lien, M. Scanziani, Nat. Neurosci. 18, 1789 (2015).

44. N. Dehghani, et al., Scientific Reports 6 (2016).

45. R. Rosenbaum, M. A. Smith, A. Kohn, J. E. Rubin, B. Doiron, Nat. Neurosci. 20, 107 (2017).

46. S. Spreizer, A. Aertsen, A. Kumar, PLOS Comput. Biol. 15, e1007432 (2019).

47. M. Diesmann, M.-O. Gewaltig, A. Aertsen, Nature 402, 529 (1999).

48. A. Litwin-Kumar, B. Doiron, Nat. Neurosci. 15, 1498 (2012).

49. B. M. Yu, et al., Journal of Neurophysiology 102, 614 (2009).

50. G. Hennequin, T. Vogels, W. Gerstner, Neuron 82, 1394 (2014).

51. G. F. Elsayed, A. H. Lara, M. T. Kaufman, M. M. Churchland, J. P. Cunningham, Nature Communications 7, 1 (2016).

52. A. Luczak, P. Barthó, K. D. Harris, Neuron 62, 413 (2009).

53. J. D. Semedo, A. Zandvakili, C. K. Machens, M. Y. Byron, A. Kohn, Neuron 102, 249 (2019).

54. M. G. Perich, et al., BioRxiv (2020).

55. A. Riehle, T. Brochier, M. Nawrot, S. Grün, Front. Neural Circuits 12 (2018).

56. P. A. Dąbrowska, et al., bioRxiv (2020).

57. P. Bartho, et al., J. Neurophysiol. 92, 600 (2004).

58. M. T. Kaufman, et al., Journal of Neurophysiology 104, 799 (2010).

59. M. T. Kaufman, M. M. Churchland, K. V. Shenoy, Journal of Neurophysiology 110, 817 (2013).

60. A. Peyrache, et al., Proc. Nat. Acad. Sci. USA 109, 1731 (2012).

61. A. Peyrache, A. Destexhe, Neurobiology of Disease 130, 104500 (2019).

62. E. Torre, et al. 36, 8329 (2016).

63. N. Hatsopoulos, J. Joshi, J.G. O’Leary, Journal of Neurophysiology 92, 1165 (2004).

64. F. W. J. Olver, D. W. Lozier, R. F. Boisvert, C. W. Clark, eds., NIST Handbook of Mathematical Functions (Cambridge University Press, New York, NY, 2010).

65. D. Grytskyy, T. Tetzlaff, M. Diesmann, M. Helias, Front. Comput. Neurosci. 7, 131 (2013).

66. C. W. Gardiner, Handbook of Stochastic Methods for Physics, Chemistry and the Natural Sciences, no. 13 in Springer Series in Synergetics (Springer-Verlag, Berlin, 1983).

67. B. Kriener, M. Helias, S. Rotter, M. Diesmann, G. T. Einevoll, Front. Comput. Neurosci. 7, 1 (2014).

68. J. Basdevant, Fortschritte der Physik 20, 283 (1972).

69. T. C. Potjans, M. Diesmann, Cereb. Cortex 24, 785 (2014).

70. J. Senk, E. Hagen, S. J. van Albada, M. Diesmann, arXiv preprint 1805.10235v1 (2018).

71. N. Goldenfeld, Lectures on phase transitions and the renormalization group (Perseus books, Reading, Mas-sachusetts, 1992).

72. M. Abramowitz, I. A. Stegun, Handbook of Mathematical Functions, vol. 55 of Applied Mathematics Series (National Bureau of Standards, Washington, 1964).

73. A. Pelizzola, Physical Review E 49, R2503 (1994).

74. J. Jordan, et al., Nest 2.18.0 (2019).

75. J. Hahne, et al. 11, 34 (2017).

76. H. C. Tuckwell, Introduction to Theoretical Neurobiology (Cambridge University Press, Cambridge, 1988), vol. 1, chap. 3, The Lapique model of the nerve cell, pp. 85–123.

77. D. J. Amit, M. V. Tsodyks, Network 2, 259 (1991).

