## Supplementary Material for "Global organization of neuronal activity only requires unstructured local connectivity"

### Appendix 1

#### 1 Robustness to E/I separation

The analysis of the experimental data involves a number of preprocessing steps, which may affect the resulting statistics. In our study one such critical step is the separation of putative excitatory and inhibitory units, which is partially based on setting thresholds on the widths of spike waveform, as described in the Methods section. We tested the robustness of our conclusions with respect to these thresholds.

As mentioned in Methods, two thresholds for the width of a spike waveform are chosen, based on all SU average waveforms: A width larger than the "broadness" threshold indicates a putative excitatory neuron. A width lower than the "narrowness" threshold indicates a putative inhibitory neuron. Units with intermediate widths are unclassified. Additionally, to increase the reliability of the classification, we perform it in two steps: first on the SU's average waveform, and second on all its single waveforms. We calculate the percentage of single waveforms classified as either type. Finally, only SUs showing a high enough percentage of single waveforms classified the same as the average waveform are sorted as the respective type. The minimal percentage required, referred to as consistency $c$, is initially set to the lowest value which ensures no contradictions between average- and single-waveform thresholding results. While the "broadness" and "narrowness" thresholds are chosen based on all available data for a given monkey, the required consistency is determined separately for each recording session. For monkey N $c$ is set to $0.6$ in all but one sessions: In resting state session N1 it is increased to $0.62$. For monkey E the values of $c$ equals $0.6$ in the resting state recordings and take the following values in five analyzed reach-to-grasp sessions: $0.6$, $0.89$, $0.65$, $0.61$, $0.64$.

The only step of our analysis for which the separation of putative excitatory and inhibitory neurons is crucial is the fitting of exponentials to the distance-resolved covariances. This step only involves resting state data. To test the robustness of our conclusions, we manipulate the required consistency value for sessions E1, E2, N1, and N2 by setting it to $0.75$. Appendix 1-figure 1 and Appendix 1-table 1 summarize the resulting fits.

It turns out that increasing $c$ to $0.75$, which implies disregarding about $20 - 25$ percent of all data, does not have a strong effect on the fitting results. The obtained decay constants are smaller than for a lower $c$ value, but they stay in a range about an order of magnitude larger than the anatomical connectivity. We furthermore see that fitting individual slopes to different populations in some sessions leads to unreliable results (cf. yellow lines in Appendix 1-figure 1a, i and blue lines in Appendix 1-figure 1c, d, k, l). Therefore, the data is not sufficient to detect differences in decay constants for different neuronal populations. Fitting instead a single decay constant yields trustworthy results (cf. yellow lines in Appendix 1-figure 1e, m and blue lines in Appendix 1-figure 1g, h, o, p). Our data thus clearly expose that decay constants of covariances are in the millimeter range.

#### 2 Stationarity of behavioral data

The linear response theory, with the aid of which we develop our predictions about the covariance structure in the network, assumes that the processes under examination are stationary in time. However, this assumption is not necessarily met in experimental data, especially in motor cortex during active behavioral tasks. For this reason we analyzed the stationarity of average single unit firing rate and pairwise zero time-lag covariance throughout a reach-to-grasp trial, similarly to (*15*). Although the spiking activity becomes highly non-stationary during the movement, those epochs that are chosen for the analysis in our study show only moderate variability in time (Appendix 1-figure 2).

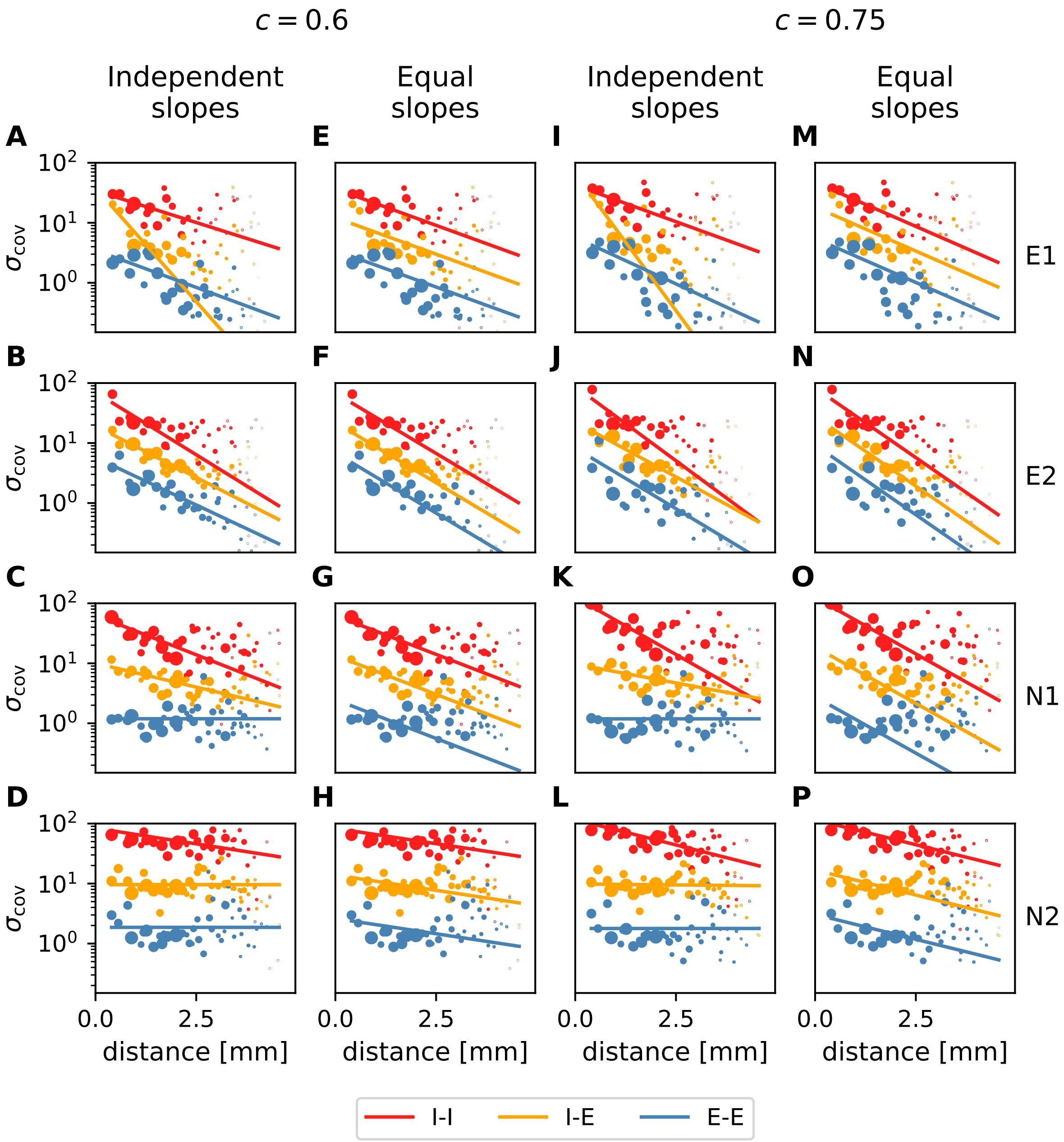


Appendix 1-figure 1: **Distance-resolved variance of covariance: robustness of decay constant estimation.** Exponential fits (lines) to variances of covariances (dots) analogous to figure 4*A*,*B* in the main text (columns 1&3 and 2&4, respectively) for all analyzed resting state sessions. The two sets of plots differ in E/I separation consistency values chosen during data preprocessing. Panels a-h: default (lowest) required consistency ($\sim 0.6$), used throughout the main analysis; panels i-p: $c = 0.75$. The values of the obtained decay constants are listed in Appendix 1-table 1. The source data and code used to create this figure are provided as supp_fig1.zip.

| $c$ | | E1 | E2 | N1 | N2 |
|---|---|---|---|---|---|
| 0.6 (default) | #exc/#inh | 56/50 | 67/56 | 76/45 | 78/62 |
| | unclassified | 0.078 | 0.075 | 0.069 | 0.091 |
| | relative error | 1.1157 | 1.0055 | 1.0097 | 1.0049 |
| | 1-slope fit | 1.674 | 1.029 | 1.676 | 4.273 |
| | I-I | 1.919 | 0.996 | 1.647 | 4.156 |
| | I-E | 0.537 | 1.206 | 2.738 | 96100.688 |
| | E-E | 1.642 | 1.308 | 80308.482 | 94096.871 |
| 0.75 | #exc/#inh | 45/42 | 47/48 | 70/36 | 74/48 |
| | unclassified | 0.24 | 0.28 | 0.18 | 0.21 |
| | relative error | 1.1778 | 1.0141 | 1.0102 | 1.0090 |
| | 1-slope fit | 1.357 | 0.874 | 1.420 | 2.587 |
| | I-I | 1.794 | 0.809 | 1.394 | 2.550 |
| | I-E | 0.496 | 1.123 | 3.682 | 40.852 |
| | E-E | 1.390 | 1.199 | 80548.500 | 10310.780 |

Appendix 1-table 1: **Summary of exponential fits to distance-resolved variance of covariance.** For each value of E/I separation consistency $c$ the numbers of sorted putative neurons and the percentages of unclassified units, and therefore not considered for fitting SUs, are listed per resting state session, along with the resulting fits (cf. figure 4 in the main text)

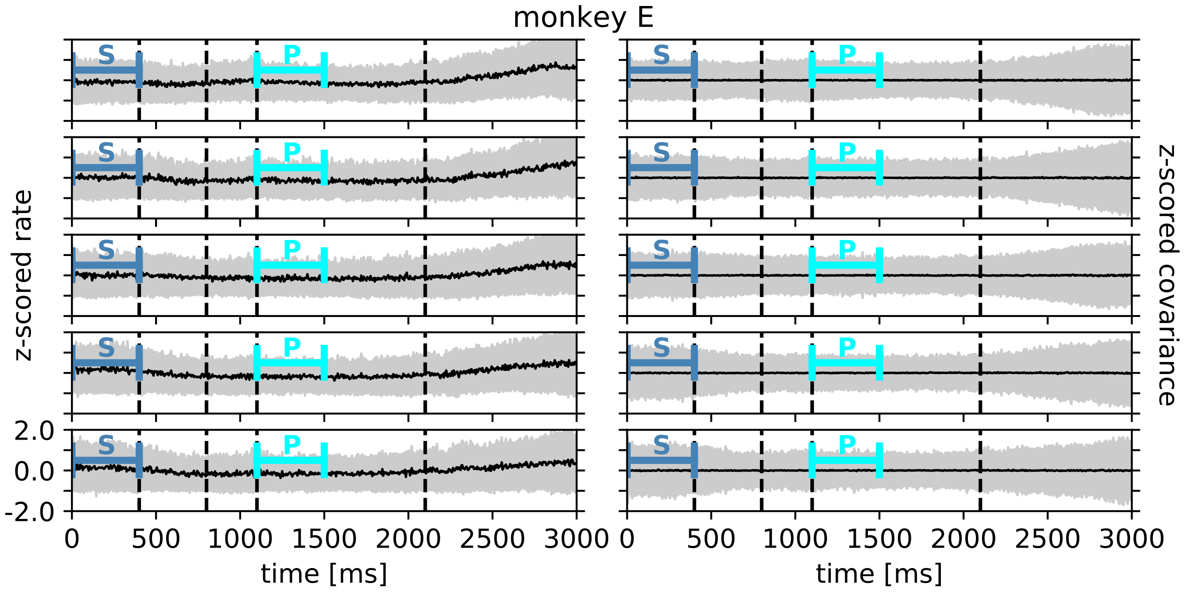


Appendix 1-figure 2: **Rate and covariance stationarity during a reach-to-grasp trial: monkey E.** Black line indicates mean and gray area +/- 1 standard deviation range of single unit firing rate (left column) and pairwise zero time-lag covariance (right column) during trial of a given session (row). Blue continuous lines indicate starting (S) and preparatory (P) periods used in the analysis (cf. figure 6 in the main text). First, second and fourth dashed lines indicate visual signals lighting up and the third dashed line indicates the removal of a visual cue and beginning of a waiting period. The source data and code used to create this figure are provided as supp_fig2.zip.

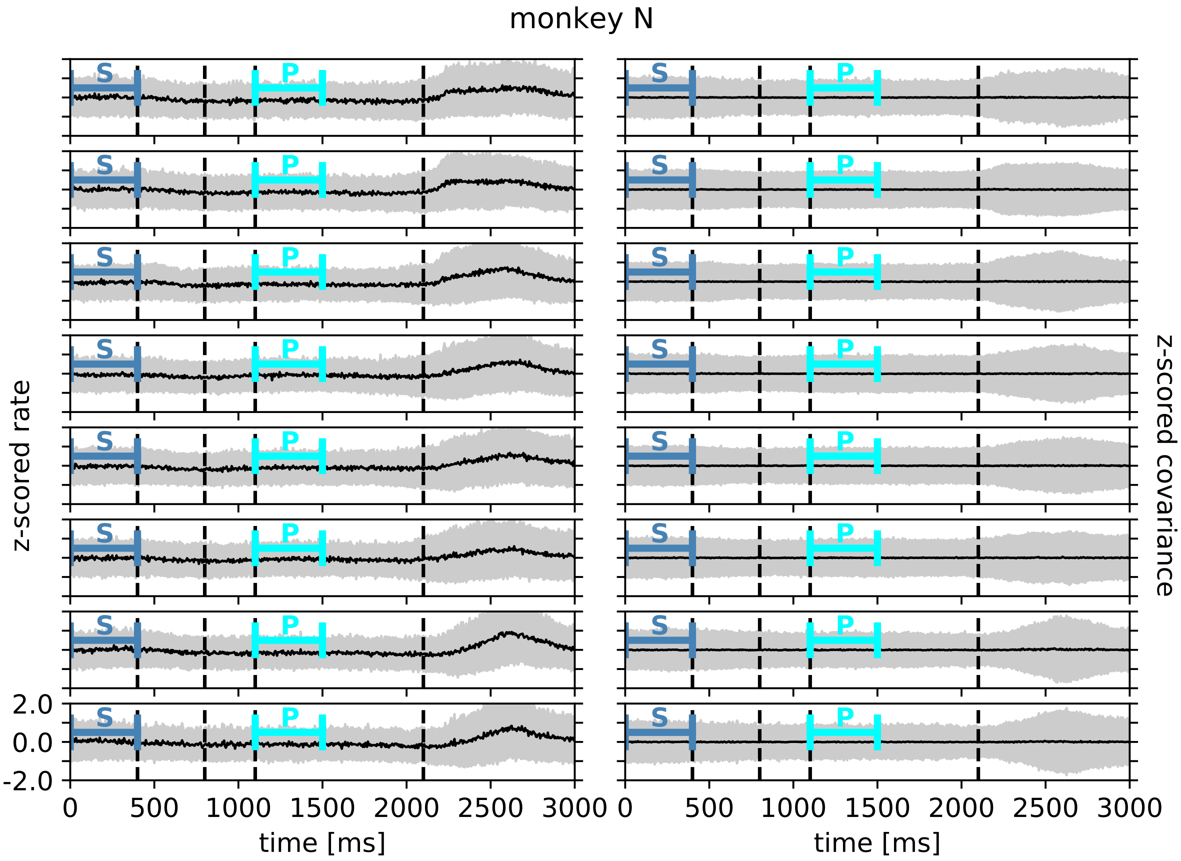


Appendix 1-figure 3: **Rate and covariance stationarity during a reach-to-grasp trial: monkey N.** Analogous to Appendix 1-figure 2. The source data and code used to create this figure are provided as supp_fig3.zip.

| session | $N_{\text{trials}}$ | $N_{\text{single units}}$ |
|---|---|---|
| e161212-002 | 108 | 129 |
| e161214-001 | 99 | 118 |
| e161222-002 | 102 | 118 |
| e170105-002 | 101 | 116 |
| e170106-001 | 100 | 113 |
| i140613-001 | 93 | 137 |
| i140617-001 | 129 | 155 |
| i140627-001 | 138 | 145 |
| i140702-001 | 157 | 134 |
| i140703-001 | 142 | 142 |
| i140704-001 | 141 | 124 |
| i140721-002 | 160 | 96 |
| i140725-002 | 151 | 106 |

Appendix 1-table 2: **Numbers of trials and single units per reach-to-grasp recording session.** Session names starting with “e” correspond to monkey E and session names starting with “i” to monkey N.

### 3 Network Model

We are considering neuronal network models with isotropic and distance-dependent connection profiles. Ultimately, we are interested in describing cortical networks with two-dimensional sheet-like structure. But, for developing the theory, we first consider the simpler case of a one-dimensional ring and subsequently develop the theory on a two-dimensional torus, ensuring periodic boundary conditions in both cases. $N$ equidistantly distributed neurons form a grid on these manifolds. The position of neuron $i \in \{1, ..., N\}$ is described by the vector $\boldsymbol{r}_i \in \mathbb{R}^D$, $D \in \{1, 2\}$. The connections $W_{ij}$ from neuron $j$ to neuron $i$ are drawn randomly with a connection probability that decays with distance between neurons $|\boldsymbol{r}_i - \boldsymbol{r}_j|$, described by the normalized connectivity profile $p(\boldsymbol{r})$, $\int p(\boldsymbol{r})\,\mathrm{d}^D r = 1$, which we assume to obey radial symmetry. The connection probability decays on a characteristic length scale $d$. As we are working on discrete lattices, we introduce the probability of two neurons being connected $p_{ij}$, which is defined by the relation $p(\boldsymbol{r}_i - \boldsymbol{r}_j) = \lim_{a\to 0} p_{ij}/a$, with lattice spacing $a$. We set the synaptic weights for connections of a single type to a fixed value $w$, but allow for multiple connections between neurons, that is $W_{ij} \in \{0, w, 2w, ...\}$ for all sending neurons $j$ of a given type.

We introduce two populations of neurons, excitatory (E) and inhibitory (I) neurons. The number of neurons of a given population $a \in \{\mathrm{E}, \mathrm{I}\}$ is $N_a$, and their ratio is $q = N_\mathrm{E}/N_\mathrm{I}$, which, for convenience, we assume to be an even number (see permutation symmetry below). The connection from population $b$ to population $a$ has the synaptic weight $w_{ab}$ and characteristic decay length of the connectivity profile $d_{ab}$. The average number of inputs drawn per neuron is fixed to $K_{ab}$. In order to preserve translation symmetry, $q$ excitatory neurons and one inhibitory neuron are put onto the same lattice point, as shown in figure 3*A* in the main text.

Linear response theory has been shown to faithfully capture the statistics of fluctuations in asynchronous irregular network states (*35*). Covariances in these states can be modeled using linear rate neurons with dynamics of their activity $\boldsymbol{x} \in \mathbb{R}^N$ described by

$$\tau \frac{\mathrm{d}}{\mathrm{d}t}\boldsymbol{x} = -\boldsymbol{x} + \boldsymbol{W}\boldsymbol{x} + \boldsymbol{\xi} \quad ,$$

with uncorrelated Gaussian white noise $\boldsymbol{\xi}$, $\langle \xi_i(t)\rangle = 0$, $\langle \xi_i(s)\xi_j(t)\rangle = D_i\delta_{ij}\delta(s-t)$. The solution to this differential equation can be found by multiplying the whole equation with the left eigenvectors $\boldsymbol{u}_\alpha$ of $\boldsymbol{W}$

$$\tau \frac{\mathrm{d}}{\mathrm{d}t} y_\alpha = -y_\alpha + \lambda_\alpha y_\alpha + \xi_\alpha \quad , \tag{5}$$

where $y_\alpha = \boldsymbol{u}_\alpha \cdot \boldsymbol{x}$, $\xi_\alpha = \boldsymbol{u}_\alpha \cdot \boldsymbol{\xi}$ and $\lambda_\alpha$ is denoting the corresponding eigenvalue of $\boldsymbol{W}$. Neglecting the noise term, the solutions are given by

$$y_\alpha(t) \propto \Theta(t) \exp\left[-\frac{t}{\tau}(1-\lambda_\alpha)\right] \quad , \tag{6}$$

with Heavyside function $\Theta(t)$. These are the eigenmodes of the linear system and they are linear combinations of the individual neuronal rates

$$y_\alpha = \sum_{i=1}^{N} (u_\alpha)_i\, x_i \quad .$$

Note that the weights $(u_\alpha)_i$ of these linear combinations depend on the details of the effective connectivity matrix $\boldsymbol{W}$. The stability of an eigenmode is determined by the corresponding eigenvalue $\lambda_\alpha$. If $\mathrm{Re}\,(\lambda_\alpha) < 1$, the eigenmode is stable and decays exponentially. If $\mathrm{Re}\,(\lambda_\alpha) > 1$, the eigenmode is unstable and grows exponentially. If $\mathrm{Im}\,(\lambda_\alpha) \neq 0$, the eigenmode is oscillatory with an exponential envelope. $\mathrm{Re}\,(\lambda_\alpha) = 1$ is here referred to as the critical point. This type of stability is also called linear stability to stress that these considerations are only valid in the linear approximation. Realistic neurons have a saturation at high rates, which prohibits divergences and therefore unstable modes do not occur in real neuronal networks. A network is called linearly stable if all modes are stable. This is determined by the real part of the largest eigenvalue of $\boldsymbol{W}$, called spectral bound $R$. In inhibition-dominated networks, the spectral bound is determined by the heterogeneity in connections and $R \lessapprox 1$ defines the dynamically balanced critical state (*15*).

The different noise components $\xi_\alpha$ excite the corresponding eigenmodes of the system and act as a driving force. A noise vector $\boldsymbol{\xi}$ that is not parallel to a single eigenvector $\boldsymbol{u}_\alpha$ excites several eigenmodes, each with the corresponding strength $\xi_\alpha$.

Note that the different eigenmodes do not interact, which is why the total activity $\boldsymbol{x}$ is given by a linear combination, or superposition, of the eigenmodes

$$\boldsymbol{x} = \sum_{\alpha=1}^{N} y_\alpha \boldsymbol{v}_\alpha \quad ,$$

where $\boldsymbol{v}_\alpha$ denotes the $\alpha-$th right eigenvector of the connectivity matrix $\boldsymbol{W}$.

### 4 Covariances

Time-lag integrated covariances $c_{ij} = \int d\tau \, \langle x_i(t) x_j(t+\tau) \rangle - \langle x_i(t) \rangle \langle x_j(t+\tau) \rangle$ can be computed analytically for the linear dynamics (5). They follow from the connectivity $\boldsymbol{W}$ and the noise strength $D$ as (*35, 24, 31, 65*)

$$\boldsymbol{c} = [\boldsymbol{1} - \boldsymbol{W}]^{-1} \boldsymbol{D} [\boldsymbol{1} - \boldsymbol{W}]^{-\mathrm{T}}, \tag{7}$$

with identity matrix $\boldsymbol{1}$. By the Wiener-Khinchin theorem, these covariances are equivalent to covariances of spike counts in large time windows (cf. figure $1B$ in the main text) that can be computed from trial-resolved spiking data (*15*). This equivalence allow us to directly relate theoretical predictions for covariances to the experimentally observed ones.

While equation (7) provides the full information on covariances between any two neurons in the network, this information is not available in the experimental data. Only a small subset of neuronal activities can be recorded such that inference of connectivity parameters from (7) is unfeasible. We recently proposed in (*15*) to instead consider the statistics of covariances as the basis for comparison between models and data. Using (5) and (7) as a starting point, field theoretical techniques allow the derivation of equations for the mean $\overline{\boldsymbol{c}}$ and variance $\overline{\delta \boldsymbol{c}^2}$ of cross-covariances in relation to the mean $\boldsymbol{M}$ and variance $\boldsymbol{S}$ of the connectivity matrix $\boldsymbol{W}$ (*15*):

$$\overline{\boldsymbol{c}} = [\boldsymbol{1} - \boldsymbol{M}]^{-1} \boldsymbol{D}_\mathrm{r} [\boldsymbol{1} - \boldsymbol{M}]^{-\mathrm{T}}, \tag{8}$$

$$\overline{\delta \boldsymbol{c}^2} = [\boldsymbol{1} - \boldsymbol{S}]^{-1} \boldsymbol{D}_\mathrm{r}^2 [\boldsymbol{1} - \boldsymbol{S}]^{-\mathrm{T}}. \tag{9}$$

$\boldsymbol{M}$ and $\boldsymbol{S}$ are defined in the subsequent section. The renormalized input noise strength is given by

$$\boldsymbol{D}_\mathrm{r} = \boldsymbol{D} + \mathrm{diag}\left[\boldsymbol{D} \left(\boldsymbol{1} - \boldsymbol{S}\right)^{-1} \boldsymbol{S} \cdot \boldsymbol{I}\right] \quad , \tag{10}$$

with input noise covariance $\boldsymbol{D}$, and the all-ones vector $\boldsymbol{I} = (1, \ldots, 1)^\mathrm{T} \in \mathbb{R}^N$. Note that Equation (9) only holds for cross-covariances ($i \neq j$). The diagonal terms $\left[\overline{\delta \boldsymbol{c}^2}\right]_{ii}$, i.e. the variance of auto-covariances, do get a second contribution, which is negligible for the cross-covariances considered here .

### 5 Cumulant Generating Function of Connectivity Matrix

For calculating the mean and variance of the covariances of the network activity ((8) and (9)) we need mean $\boldsymbol{M}$ and variance $\boldsymbol{S}$ of connectivity $\boldsymbol{W}$. In the following, we derive the cumulant generating function (*66*) of $W_{ij}$.

We assume that connections for each neuron pair are drawn independently. The number of connections $n$ from neuron $j$ to neuron $i$ is a Binomial random variable with $K$ trials with the probability of success given by $p_{ij}$ (in the following, for brevity, we ignore the index $i$, $p_{ij} \equiv p_j$)

$$\mathrm{Prob}_j(n) = \binom{K}{n} p_j^n \left(1 - p_j\right)^{K-n} \quad .$$

The average number of connections from neuron $j$ to neuron $i$ is $K_j = p_j K$, which assures the correct average total indegree

$$\sum_j K_j = K \sum_j p_j = K \quad .$$

The moment generating function of a connectivity matrix element $W_j \equiv W_{ij} \in \{0, w, 2w, ...\}$ is given by

$$Z_{W_j}(k) = \sum_{n=0}^{K} \binom{K}{n} p_j^n \left(1 - p_j\right)^{K-n} \mathrm{e}^{nwk} \quad .$$

In a realistic network, $K$ is very large. In the limit $K \to \infty$, while keeping $Kp = \mathrm{const.}$, the Binomial distribution converges to a Poisson distribution and we can write

$$\begin{aligned} Z_{W_j}(k) &\approx \sum_{n=0}^{K} \frac{K_j^n}{n!} \mathrm{e}^{-K_j} \mathrm{e}^{nwk} \\ &= \sum_{n=0}^{K} \frac{\left(K_j \mathrm{e}^{wk}\right)^n}{n!} \mathrm{e}^{-K_j} \\ &\xrightarrow{K \to \infty} \exp\left[K_j \left(\mathrm{e}^{wk} - 1\right)\right] \quad . \end{aligned}$$

Taking the logarithm leads to the cumulant generating function

$$G_{W_j}(k) \approx p_j K \left(\mathrm{e}^{wk} - 1\right) \quad ,$$

and the first two cumulants

$$\begin{aligned} M_{ij} &= \frac{\partial}{\partial k} G_{W_j}(k) \bigg|_{k=0} = p_j K w = p\left(|x_i - x_j|\right) K w \quad , \\ S_{ij} &= \frac{\partial^2}{\partial k^2} G_{W_j}(k) \bigg|_{k=0} = p_j K w^2 = p\left(|x_i - x_j|\right) K w^2 \quad . \end{aligned}$$

### 6 Note on Derivation of Variance of Covariances

Note that $\boldsymbol{M}$ and $\boldsymbol{S}$ have an identical structure determined by the connectivity profile and the structure of the covariance equation is identical for the mean Equation (8) and variance Equation (9) as well. This is why in the following we only derive the results for the mean of covariances. The results for the variance of covariances is obtained by substituting $w$ by $w^2$ and $\boldsymbol{D}_\mathrm{r}$ by $\boldsymbol{D}_\mathrm{r}^2$. As we show, divergences in expressions related to the mean covariances arise if the population eigenvalue $\lambda_0$ of the effective connectivity matrix approaches one. In expressions related to the variance of covariances, the divergences are caused by the squared spectral bound $R^2$ being close to one. In general expressions, we sometimes write $\zeta$ in order to denote either the population eigenvalue or the spectral bound, corresponding to the context of mean or variance of covariances.

### 7 Utilizing Symmetries to Reduce Dimensionality

For real neuronal networks, the anatomical connectivity is never known completely, let alone the effective connectivity. This is why we are considering disorder-averaged systems. They are described by the mean $\boldsymbol{M}$ and variance $\boldsymbol{S}$ of the connectivity. The latter inherit the underlying symmetries of the network, like for example the same radially symmetric connectivity profile for all neurons of one type. As neuronal networks are high dimensional systems, calculating covariances from Equation (8) and Equation (9) first seems like a daunting task. But, leveraging the aforementioned symmetries similarly as in (*67*) allows for an effective reduction of the dimensionality of the system, thereby rendering the problem manageable.

As a demonstrative example of how this is done, consider a random network of $N$ neurons on a one-dimensional ring, in which a neuron can form a connection with weight $w$ to any other neuron with probability $p_0$. In that case, $\boldsymbol{M}$ is a homogeneous matrix, with all entries given by the same average connectivity weight

$$\boldsymbol{M} = \begin{pmatrix} p_0 w & p_0 w & \dots & p_0 w \\ p_0 w & p_0 w & \dots & p_0 w \\ \vdots & \vdots & \ddots & \vdots \\ p_0 w & p_0 w & \dots & p_0 w \end{pmatrix} \quad .$$

This corresponds to an all-to-all connected ring network. Due to the symmetry of the system, moving all neurons by one lattice constant does not change the system. The translation operator $\boldsymbol{T}$, representing this operation mathematically, is defined via its effect on the vector of neuron activity $\boldsymbol{x}$

$$\boldsymbol{T}\boldsymbol{x} = \boldsymbol{T} \begin{pmatrix} x_1 \\ x_2 \\ \vdots \\ x_N \end{pmatrix} = \boldsymbol{T} \begin{pmatrix} x_N \\ x_1 \\ \vdots \\ x_{N-1} \end{pmatrix} \quad .$$

Applying $\boldsymbol{T}$ $N$-times yields the identity operation

$$\boldsymbol{T}^N = \mathbf{1} \quad .$$

Hence, its eigenvalues are given by complex roots of one

$$\mathrm{e}^{-\mathrm{i}2\pi l/N} = \mathrm{e}^{-\mathrm{i}2\pi la/L} = \mathrm{e}^{-\mathrm{i}k_l a} \quad , \quad l \in \{0, 1, ..., N-1\} \quad ,$$

with $L = Na$ denoting the circumference of the ring. This shows that $\boldsymbol{T}$ has $N$ one-dimensional eigenspaces. Since the system is invariant under translation, $\boldsymbol{M}$ is invariant under the transformation

$$\boldsymbol{T}\boldsymbol{M}\boldsymbol{T}^{-1} = \boldsymbol{M} \quad ,$$

and thus $\boldsymbol{M}$ and $\boldsymbol{T}$ commute. As $\boldsymbol{M}$ leaves eigenspaces of $\boldsymbol{T}$ invariant (if $\boldsymbol{v}$ is an eigenvector of $\boldsymbol{T}$, $\boldsymbol{M}\boldsymbol{v}$ is an eigenvector with the same eigenvalue, so they need to be multiples of each other), all eigenvectors of $\boldsymbol{T}$ must be eigenvectors of $\boldsymbol{M}$. Accordingly, knowing the eigenvectors of $\boldsymbol{T}$ allows diagonalizing $\boldsymbol{M}$. The normalized (left and right) eigenvectors of $\boldsymbol{T}$ are given by

$$\boldsymbol{v}_{k_l} = \frac{1}{\sqrt{N}} \begin{pmatrix} 1 \\ \mathrm{e}^{\mathrm{i}k_l a} \\ \mathrm{e}^{\mathrm{i}2k_l a} \\ \vdots \\ \mathrm{e}^{\mathrm{i}(N-1)k_l a} \end{pmatrix} \quad .$$

We get the eigenvalues of $\boldsymbol{M}$ by multiplying it with the eigenvectors of $\boldsymbol{T}$

$$\begin{aligned} m_{k_l} &= \boldsymbol{v}_{k_l}^\dagger \boldsymbol{M} \boldsymbol{v}_{k_l} \\ &= \frac{1}{N} p_0 w \boldsymbol{v}_{k_l}^\dagger \cdot \begin{pmatrix} \sum_{j=0}^{N-1} \mathrm{e}^{\mathrm{i}k_l aj} \\ \sum_{j=0}^{N-1} \mathrm{e}^{\mathrm{i}k_l aj} \\ \vdots \\ \sum_{j=0}^{N-1} \mathrm{e}^{\mathrm{i}k_l aj} \end{pmatrix} \quad , \end{aligned}$$

which is always zero, except for $l = 0$, which corresponds to the population eigenvalue $\lambda_0 := m_{k_0} = Np_0 w$ of $\boldsymbol{W}$ (figure 3$C$ in the main text). Now, we can simply write down the diagonalized form of $\boldsymbol{M}$

$$\begin{pmatrix} \lambda_0 & 0 & \dots & 0 \\ 0 & 0 & \dots & 0 \\ \vdots & \vdots & \ddots & \vdots \\ 0 & 0 & \dots & 0 \end{pmatrix} \quad ,$$

and we effectively reduced the $N$-dimensional to a one dimensional problem. Inverting $\boldsymbol{A} := \boldsymbol{1} - \boldsymbol{M}$ in Equation (8) is straightforward now, since it is diagonal in the new basis. Its eigenvalues can be written as

$$a_k = 1 - m_k \quad ,$$

where we suppressed the index $l$. Therefore its inverse is given by

$$\begin{aligned} A_{ij}^{-1} &= \sum_k a_k^{-1} \left(\boldsymbol{v}_k\right)_i \left(\boldsymbol{v}_k^\dagger\right)_j \\ &= \frac{1}{N} \sum_k \frac{1}{1 - m_k} \mathrm{e}^{\mathrm{i}k(x_i - x_j)} \\ &= \frac{1}{N} \sum_k \left(1 + \frac{m_k}{1 - m_k}\right) \mathrm{e}^{\mathrm{i}k(x_i - x_j)} \\ &= \delta_{ij} + \frac{1}{N} \frac{\lambda_0}{1 - \lambda_0}. \end{aligned}$$

The renormalized noise can be evaluated using that the all-ones vector occurring in equation Equation (10) is the eigenvector $\boldsymbol{v}_0$ of $\boldsymbol{S}$. After identifying the eigenvalue $s_0$ with the squared spectral bound $R^2$, we find

$$\boldsymbol{D}_\mathrm{r} = \mathrm{diag}\left(\frac{D}{1 - R^2}\right) \quad ,$$

which allows us to express the mean cross-covariances $\overline{c}$ (see Equation (8)) and the variance of cross-covariances $\overline{\delta c^2}$ (see Equation (9)) in terms of the eigenvectors of $\boldsymbol{M}$ and $\boldsymbol{S}$ respectively

$$\overline{c} = \frac{D}{1 - R^2} \left\{ \frac{2\lambda_0}{N\left(1 - \lambda_0\right)} + \left[\frac{\lambda_0}{N\left(1 - \lambda_0\right)}\right]^2 N \right\} \quad ,$$

$$\overline{\delta c^2} = \left(\frac{D}{1 - R^2}\right)^2 \left\{ \frac{2R^2}{N\left(1 - R^2\right)} + \left[\frac{R^2}{N\left(1 - R^2\right)}\right]^2 N \right\} \quad .$$

### 8 One-Dimensional Network with One Population

The simplest network with spatial connectivity is a one-dimensional ring of neurons with one population of neurons. Following section Section 5, the mean connectivity matrix has the form

$$\boldsymbol{M} = Kw \begin{pmatrix} p_{11} & p_{12} & \dots & p_{1N} \\ p_{21} & p_{22} & \dots & p_{2N} \\ \vdots & \vdots & \ddots & \vdots \\ p_{N1} & p_{N2} & \dots & p_{NN} \end{pmatrix} \quad .$$

As $p_{ij}$ only depends on the distance of two neurons, the rows in $\boldsymbol{M}$ are identical, but shifted by one index.

#### 8.1 Dimensionality Reduction

We follow the procedure developed in Section 7, as the system is invariant under translation as well. Suppressing the subscripts of $k$, we get the eigenvalues of $\boldsymbol{M}$

$$
\begin{aligned}
m_k &= \frac{1}{N} K w \left(1, \mathrm{e}^{-\mathrm{i}ka}, ..., \mathrm{e}^{-\mathrm{i}(N-1)ka}\right) \begin{pmatrix} \sum_{j=0}^{N-1} p_{1(j+1)} \mathrm{e}^{\mathrm{i}kaj} \\ \sum_{j=0}^{N-1} p_{2(j+1)} \mathrm{e}^{\mathrm{i}kaj} \\ \vdots \\ \sum_{j=0}^{N-1} p_{N(j+1)} \mathrm{e}^{\mathrm{i}kaj} \end{pmatrix} \\
&= K w \sum_{j=0}^{N-1} p_{(j+1)} \mathrm{e}^{\mathrm{i}kaj} \\
&= K w a \sum_{x} p(x) \mathrm{e}^{-\mathrm{i}kx} \quad ,
\end{aligned}
$$

where the sum over $\boldsymbol{x}$ denotes a sum over all lattice sites. We used the translational symmetry from the first to the second line. The change of sign in the exponential from line two to three is due to the fact that we are summing over the second index of $p_{ij}$. Thus, the eigenvalues are effectively given by the discrete Fourier transform of the connectivity profile. Expressing $\boldsymbol{A}^{-1}$ using the eigenvectors $\boldsymbol{v}_k$ of $\boldsymbol{M}$ leads to

$$
\begin{aligned}
A_{ij}^{-1} &= \frac{1}{N} \sum_k \frac{1}{1-m_k} \mathrm{e}^{\mathrm{i}k(x_i - x_j)} \\
&= \frac{1}{N} \sum_k \left(1 + \frac{m_k}{1-m_k}\right) \mathrm{e}^{\mathrm{i}k(x_i - x_j)} \\
&= \delta_{ij} + \frac{1}{N} \sum_k \frac{m_k}{1-m_k} \mathrm{e}^{\mathrm{i}k(x_i - x_j)} \\
&\equiv \delta_{ij} + \mu_{ij} \quad ,
\end{aligned}
\tag{11}
$$

where we extracted an identity for later convenience, and we defined $\mu_{ij}$.

Next, we consider the renormalized noise, which is given by Equation (10). Using that the all-ones vector $\boldsymbol{I}$ in the second term is the eigenvector of $\boldsymbol{S}$ corresponding to $k = 0$, we get

$$
\boldsymbol{D} \left(\boldsymbol{1} - \boldsymbol{S}\right)^{-1} \boldsymbol{S} \cdot \boldsymbol{v_0} = D \frac{s_0}{1 - s_0} \quad .
$$

Again, we identify $s_0$ with the spectral bound $R^2$, and find

$$
D_{\mathrm{r}} = D + D \frac{R^2}{1-R^2} = \frac{D}{1-R^2} \quad . \tag{12}
$$

Inserting Equation (11) and Equation (12) into Equation (8) yields

$$
\bar{c}_{ij} = \frac{D}{1-R^2} \left(\delta_{ij} + 2\mu_{ij} + \sum_k \mu_{ik} \mu_{kj}\right) \quad .
$$

#### 8.2 Continuum Limit

As we assume the lattice constant to be small, we know that the connectivity profile is sampled densely, and we are allowed to take the continuum limit. Therefore, we write

$$\begin{aligned} m_k &= Kw \sum_j p_{(j+1)} \mathrm{e}^{\mathrm{i}kaj} \\ &= Kw \sum_j a \frac{p_{(j+1)}}{a} \mathrm{e}^{\mathrm{i}kaj} \\ &\xrightarrow{a \to 0} Kw \int_{-L/2}^{L/2} \mathrm{d}x \, p(-x) \mathrm{e}^{\mathrm{i}kx} \\ &= Kw \int_{-L/2}^{L/2} \mathrm{d}x \, p(x) \mathrm{e}^{-\mathrm{i}kx} \quad . \end{aligned}$$

Note that $\lim_{a\to 0} \sum_j p_j / a = \lim_{a \to 0} \sum_j p(x_i - x_j)/a = \int dx \, p(-x)$, because we are summing over the second index $j$. If the decay constant $d$ of the connectivity profile is small compared to the size of the network $L$, we can take $L$ to infinity and finally end up with

$$m(k) = Kw \int \mathrm{d}x \, p(x) \mathrm{e}^{-\mathrm{i}kx} \quad . \tag{13}$$

Analogously, we find

$$A^{-1}(x) = \delta(x) + \frac{1}{2\pi} \int \mathrm{d}k \, \frac{m(k)}{1 - m(k)} \mathrm{e}^{\mathrm{i}kx} \equiv \delta(x) + \mu(x) \quad , \tag{14}$$

where we defined

$$\mu(x) = \frac{1}{2\pi} \int \mathrm{d}k \, \mu(k) \mathrm{e}^{\mathrm{i}kx} \quad , \tag{15}$$

with

$$\mu(k) = \frac{m(k)}{1 - m(k)} \quad . \tag{16}$$

Finally, we get

$$\bar{c}(x) = \frac{D}{1 - R^2} \left[ \delta(x) + 2\mu(x) + (\mu * \mu)(x) \right] \quad , \tag{17}$$

where the asterisk denotes the convolution.

#### 8.3 Prediction of Exponential Decay of Covariance Statistics

Note that the integral in equation Equation (15) can be interpreted as an integral in the complex plane. According to the residue theorem, the solution to this integral is a weighted sum of exponentials, evaluated at the poles of $[1 - m(k)]^{-1}$. As $\mu(x)$ appears in the equation for the mean covariances, and the convolution of two exponentials is an exponential with the prefactor (const. + |x|), we expect the dominant behavior to be an exponential decay in the long-range limit, with decay constants given by the inverse imaginary part of the poles. The poles which are closest to zero are the ones which lead to the most shallow and thereby dominant decay. A real part of the poles leads to oscillations in $\mu(x)$.

#### 8.4 Long-Range Limit

We cannot expect to solve the integral in Equation (14) for arbitrary connectivity profiles. To continue our analysis, we make use of the Padé method, which approximates arbitrary functions as rational functions (*68*). We approximate $\mu(k)$ around $k = 0$ using a Padé approximation of order (0,2)

$$\mu(k) \approx \frac{m(0)}{1 - m(0) - \frac{m''(0)}{2m(0)}k^2} \quad ,$$

with

$$m(0) = Kw \int \mathrm{d}x\, p(x) = Kw = \lambda_0 \quad , \tag{18}$$
$$m''(0) = -Kw \int \mathrm{d}x\, x^2 p(x) = -Kw \left\langle x^2 \right\rangle \quad .$$

This allows us to calculate the approximate poles of $\mu(k)$

$$k_0 = \pm\sqrt{\frac{2m(0)}{m''(0)} \left[1 - m(0)\right]} \quad . \tag{19}$$

As $2m(0)/m''(0)$ will be negative, due to factor $i^2$ from the second derivative of the Fourier integral, we write

$$k_0 = \pm \mathrm{i} \sqrt{-\frac{2m(0)}{m''(0)} \left[1 - m(0)\right]} \quad .$$

Closing the integral contour in Equation (15) in the upper half plane for $x > 0$, and in the lower half plane for $x < 0$, we get

$$\mu(x) = -\frac{m(0)^2}{m''(0)} \sqrt{-\frac{m''(0)}{2m(0)} \frac{1}{1 - m(0)}} \exp\left(-\frac{|x|}{\sqrt{-\frac{m''(0)}{2m(0)}\frac{1}{1-m(0)}}}\right) \equiv -\frac{m(0)^2}{m''(0)} d_{\mathrm{eff},\mu} \exp\left(-\frac{|x|}{d_{\mathrm{eff},\mu}}\right) \quad ,$$

where we defined the effective decay constant for the mean covariances

$$\bar{d} = \sqrt{-\frac{m''(0)}{2m(0)} \frac{1}{1 - m(0)}} = \sqrt{\frac{\langle x^2 \rangle}{2} \frac{1}{1 - \lambda_0}} \quad ,$$

with $m(0) = \lambda_0$ and $m''(0) = \lambda_0 \langle x^2 \rangle$, since $m(k)$ is the Fourier transform of the connectivity profile Equation (13). Note that $\lambda_0 = Kw$ again is the population eigenvalue of the effective connectivity matrix $\boldsymbol{W}$. For evaluating Equation (8) and Equation (9), we need to calculate the convolution of $\mu$ with itself

$$\left(\mu * \mu\right)(x) = \int \mathrm{d}y\, \mu(x - y)\mu(y) = \frac{m(0)^4}{m''(0)^2} \bar{d}^2 \left(\bar{d} + |x|\right) \exp\left(-\frac{|x|}{\bar{d}}\right) \quad .$$

The final expression for the mean covariances is

$$\bar{c}(x) = \frac{D}{1 - R^2} \left\{ \delta(x) + \left[ \left( \frac{m(0)^4}{m''(0)^2} \bar{d}^2 - 2\frac{m(0)^2}{m''(0)} \right) \bar{d} + \frac{m(0)^4}{m''(0)^2} \bar{d}^2 \, |x| \right] \exp\left(-\frac{|x|}{\bar{d}}\right) \right\} \quad .$$

Equivalently, for the variance of covariances we obtain the final result

$$\overline{\delta c^2}(x) = \left(\frac{D}{1-R^2}\right)^2 \left\{\delta(x) + \left[\left(\frac{s(0)^4}{s''(0)^2} d_{\text{eff}}^2 - 2\frac{s(0)^2}{s''(0)}\right) d_{\text{eff}} + \frac{s(0)^4}{s''(0)^2} d_{\text{eff}}^2 \, |x|\right] \exp\left(-\frac{|x|}{d_{\text{eff}}}\right)\right\} \quad ,$$

where

$$s(k) = Kw^2 \int \mathrm{d}x \, p(x) \mathrm{e}^{-\mathrm{i}kx} \quad .$$

Note that the quality of the Padé approximation depends on the outlier eigenvalue and the spectral bound. For the variances, the approximation works best for spectral bounds $R$ close to 1. The reason for this is that we are approximating the position of the poles in the complex integral Equation (15). We make an approximation around $k = 0$ and Equation (19) shows that the position of the complex poles moves closer to $k = 0$ as $s(0) \equiv R^2 \to 1$.

**General Results:** Using Equation (18)

$$m(0) = Kw = \lambda_0 \quad , \quad m''(0) = -Kw \left\langle x^2 \right\rangle \quad ,$$

we find

$$\overline{c}(x) = \frac{D}{1-R^2} \left\{\delta(x) + \left[\frac{Kw\,(1-3Kw)}{2\left\langle x^2\right\rangle(1-Kw)} \overline{d} + \frac{(Kw)^2}{\left\langle x^2\right\rangle^2} \overline{d}^2 \, |x|\right] \exp\left(-\frac{|x|}{\overline{d}}\right)\right\} \quad ,$$

with

$$\overline{d} = \sqrt{\left|\frac{\langle x^2\rangle}{2} \frac{1}{1-\lambda_0}\right|} \quad .$$

For the variance we use

$$s(0) = Kw^2 = R^2 \quad , \quad s''(0) = -Kw^2 \left\langle x^2 \right\rangle \quad ,$$

to get

$$\overline{\delta c^2}(x) = \frac{D^2}{(1-R^2)^2} \left\{\delta(x) + \left[\frac{Kw^2\left(1-3Kw^2\right)}{2\left\langle x^2\right\rangle\left(1-Kw^2\right)} d_{\text{eff}} + \frac{\left(Kw^2\right)^2}{\left\langle x^2\right\rangle^2} d_{\text{eff}}^2 \, |x|\right] \exp\left(-\frac{|x|}{d_{\text{eff}}}\right)\right\} \quad ,$$

with

$$d_{\text{eff}} = \sqrt{\left|\frac{\langle x^2\rangle}{2} \frac{1}{1-R^2}\right|} \quad .$$

**Exponential Connectivity Profile:** Using an exponential connectivity profile given by

$$p(x) = \frac{1}{2d} \mathrm{e}^{-|x|/d} \quad ,$$

we find $\left\langle x^2 \right\rangle = 2d^2$ and

$$\overline{d} = \sqrt{\left|\frac{1}{1-\lambda_0}\right|} d, \quad d_{\text{eff}} = \sqrt{\left|\frac{1}{1-R^2}\right|} d \quad ,$$

with $\lambda_0 = Kw$ for the mean, and $R^2 = Kw^2$ for the variance.

**Gaussian Connectivity Profile:** Analogously, using a Gaussian connectivity profile given by

$$p(x) = \frac{1}{\sqrt{2\pi d^2}} \mathrm{e}^{-\mathrm{x}^2/(2\mathrm{d}^2)} \quad ,$$

we find $\langle x^2 \rangle = d^2$, and get

$$\overline{d} = \sqrt{\left|\frac{1}{2}\frac{1}{1-\lambda_0}\right|} d, \quad d_{\mathrm{eff}} = \sqrt{\left|\frac{1}{2}\frac{1}{1-R^2}\right|} d \quad . \tag{20}$$

### 9 One-Dimensional Network with Two Populations

Realistic neuronal network consist of excitatory and inhibitory neurons. So we need to introduce a second population to our network. Typically, there are more excitatory than inhibitory neurons in the brain. Therefore, we introduce $q$ excitatory neurons for each inhibitory neuron. We place $q$ excitatory neurons and one inhibitory neuron together in one cell. The cells are distributed equally along the ring. For convenience, we define $N \equiv N_{\mathrm{I}}$.

The structure of the connectivity matrix depends on the choice of the activity vector $\boldsymbol{x}$. For later convenience we choose

$$\boldsymbol{x} = \begin{pmatrix} \boldsymbol{x}_1^{(\mathrm{E})} \\ x_1^{(\mathrm{I})} \\ \boldsymbol{x}_2^{(\mathrm{E})} \\ x_2^{(\mathrm{I})} \\ \vdots \\ \boldsymbol{x}_N^{(\mathrm{E})} \\ x_N^{(\mathrm{I})} \end{pmatrix} \quad ,$$

where $\boldsymbol{x}_i^{(\mathrm{E})}$ is a $q$-dimensional vector denoting the activity of the $q$ excitatory neurons in cell $i$. $\boldsymbol{M}$ is a $(q+1)N \times (q+1)N$-matrix, which qualitatively has the structure

$$\boldsymbol{M} = \begin{pmatrix} \mathrm{EE}_{11} & \mathrm{EI}_{11} & \mathrm{EE}_{12} & \mathrm{EI}_{12} & \cdots & \mathrm{EE}_{1N} & \mathrm{EI}_{1N} \\ \mathrm{IE}_{11} & \mathrm{II}_{11} & \mathrm{IE}_{12} & \mathrm{II}_{12} & \cdots & \mathrm{IE}_{1N} & \mathrm{II}_{1N} \\ \mathrm{EE}_{21} & \mathrm{EI}_{21} & \mathrm{EE}_{22} & \mathrm{EI}_{22} & \cdots & \mathrm{EE}_{2N} & \mathrm{EI}_{2N} \\ \mathrm{IE}_{21} & \mathrm{II}_{21} & \mathrm{IE}_{22} & \mathrm{II}_{22} & \cdots & \mathrm{IE}_{2N} & \mathrm{II}_{2N} \\ \vdots & \vdots & \vdots & \vdots & \ddots & \vdots & \vdots \\ \mathrm{EE}_{N1} & \mathrm{EI}_{N1} & \mathrm{EE}_{N2} & \mathrm{EI}_{N2} & \cdots & \mathrm{EE}_{NN} & \mathrm{EI}_{NN} \\ \mathrm{IE}_{N1} & \mathrm{II}_{N1} & \mathrm{IE}_{N2} & \mathrm{II}_{N2} & \cdots & \mathrm{IE}_{NN} & \mathrm{II}_{NN} \end{pmatrix} \quad . \tag{21}$$

Note that $\mathrm{EE}_{ij}$ are $q \times q$ matrices, $\mathrm{EI}_{ij}$ are $q \times 1$ matrices, $\mathrm{IE}_{ij}$ are $1 \times q$ matrices and $\mathrm{II}_{ij}$ are $1 \times 1$ matrices. The entries $ab_{ij}$ describe the connectivities from population $b$ in cell $j$ to population $a$ in cell $i$. The entries are given by

$$ab_{ij} = \begin{cases} \frac{1}{q} w_{ab} K_{ab} \left(p_{ab}\right)_{ij} & \text{if } b = \mathrm{E} \\ w_{ab} K_{ab} \left(p_{ab}\right)_{ij} & \text{if } b = \mathrm{I} \end{cases} \quad .$$

The difference stems from the fact that we have $q$ times as many excitatory neurons. As the total number of indegrees from excitatory neurons should be given by $K_{aE}$, we need to introduce a reducing factor of $1/q$, as the connection probability is normalized to one.

#### 9.1 Dimensionality Reduction

In the following, we will reduce the dimensionality of $\boldsymbol{M}$ as done before in the case with one population. First, we make use of the symmetry within the cells. All entries in $\boldsymbol{M}$ corresponding to connections coming from excitatory neurons of the same cell need to be the same. For that reason, we change the basis to

$$e_i^{(\mathrm{E})} = \frac{1}{\sqrt{q}} \begin{pmatrix} 0 \\ 0 \\ \vdots \\ \boldsymbol{I} \\ \vdots \\ 0 \end{pmatrix} \quad , \quad e_i^{(\mathrm{I})} = \begin{pmatrix} 0 \\ 0 \\ \vdots \\ 1 \\ \vdots \\ 0 \end{pmatrix} \quad , \tag{22}$$

where $\boldsymbol{I}$ denotes a $q$-dimensional vector containing only ones. For a full basis, we need to include all the vectors with $\boldsymbol{I}$ being replaced by a vector containing all possible permutations of equal numbers of $\pm 1$. In this basis $\boldsymbol{M}$ is block diagonal

$$\begin{pmatrix} \boldsymbol{M}' & 0 \\ 0 & 0 \end{pmatrix} \quad ,$$

and $\boldsymbol{M}'$ is an $2N \times 2N$ matrix, which has the same qualitative structure as shown in Equation (21), but the submatrices $(\mathrm{ab})_{ij}$ are replaced by

$$ab_{ij} = \begin{cases} w_{\mathrm{EE}} K_{\mathrm{EE}} \left(p_{\mathrm{EE}}\right)_{ij} & \text{if } ab = \mathrm{EE} \\ \sqrt{q} w_{\mathrm{EI}} K_{\mathrm{EI}} \left(p_{\mathrm{EI}}\right)_{ij} & \text{if } ab = \mathrm{EI} \\ w_{\mathrm{IE}} K_{\mathrm{IE}} \left(p_{\mathrm{IE}}\right)_{ij} / \sqrt{q} & \text{if } ab = \mathrm{IE} \\ w_{\mathrm{II}} K_{\mathrm{II}} \left(p_{\mathrm{II}}\right)_{ij} & \text{if } ab = \mathrm{II} \end{cases} \quad .$$

Next, we use translational symmetry of the cells. The translation operator is defined by

$$\boldsymbol{T}\boldsymbol{x} = \boldsymbol{T} \begin{pmatrix} \boldsymbol{x}_1^{(\mathrm{E})} \\ x_1^{(\mathrm{I})} \\ \boldsymbol{x}_2^{(\mathrm{E})} \\ x_2^{(\mathrm{I})} \\ \vdots \\ \boldsymbol{x}_N^{(\mathrm{E})} \\ x_N^{(\mathrm{I})} \end{pmatrix} = \begin{pmatrix} \boldsymbol{x}_N^{(\mathrm{E})} \\ x_N^{(\mathrm{I})} \\ \boldsymbol{x}_1^{(\mathrm{E})} \\ x_1^{(\mathrm{I})} \\ \vdots \\ \boldsymbol{x}_{N-1}^{(\mathrm{E})} \\ x_{N-1}^{(\mathrm{I})} \end{pmatrix} \quad .$$

As the system is invariant under moving each cell to the next lattice site, $\boldsymbol{M}'$ is invariant under the transformation

$$\boldsymbol{T}\boldsymbol{M}'\boldsymbol{T}^{-1} = \boldsymbol{M}' \quad .$$

Again, the eigenvalues of $\boldsymbol{T}$ can be determined using $\boldsymbol{T}^N = 1$ and they are the same as in the case of one population. But, note that here the eigenspaces corresponding to the single eigenvalues are two dimensional. The eigenvectors

$$\boldsymbol{v}_k^{(\mathrm{E})} = \frac{1}{\sqrt{N}} \begin{pmatrix} 1 \\ 0 \\ e^{\mathrm{i}ka} \\ 0 \\ \vdots \\ e^{\mathrm{i}(N-1)ka} \\ 0 \end{pmatrix} \quad , \quad \boldsymbol{v}_k^{(\mathrm{I})} = \frac{1}{\sqrt{N}} \begin{pmatrix} 0 \\ 1 \\ 0 \\ e^{\mathrm{i}ka} \\ \vdots \\ 0 \\ e^{\mathrm{i}(N-1)ka} \end{pmatrix} \quad ,$$

belong to the same eigenvalue. In this basis, $\boldsymbol{M}'$ is block diagonal, with each block consisting of a $2 \times 2$ matrix, corresponding to one value of $k_l = \frac{2\pi l}{L}$, $l \in \{0, ..., N-1\}$

$$\boldsymbol{M}' = \begin{pmatrix} \boldsymbol{M}_{k_0} & 0 & \cdots & 0 \\ 0 & \boldsymbol{M}_{k_1} & \cdots & 0 \\ \vdots & \vdots & \ddots & \vdots \\ 0 & 0 & \cdots & \boldsymbol{M}_{k_{N-1}} \end{pmatrix} \quad .$$

Since all block matrices can be treated equally, we further reduced the problem to diagonalizing a $2 \times 2$ matrix. The submatrices take the form

$$\boldsymbol{M}_k = \begin{pmatrix} m_{\mathrm{EE}}(k) & \sqrt{q}\, m_{\mathrm{EI}}(k) \\ m_{\mathrm{IE}}(k)/\sqrt{q} & m_{\mathrm{II}}(k) \end{pmatrix} \quad ,$$

with the discrete Fourier transform

$$m_{ab}(k) = K_{ab} w_{ab} \sum_{x=-Na/2}^{Na/2} p_{ab}(x) \mathrm{e}^{-ikx} \quad . \tag{23}$$

Note that $x$ and $k$ are still discrete here, but we could take the continuum limit at this point. The eigenvalues of $\boldsymbol{M}_k$ are given by

$$m_{\pm}(k) = \frac{1}{2}(m_{\mathrm{EE}}(k) + m_{\mathrm{II}}(k)) \pm \frac{1}{2}\sqrt{m_{\mathrm{EE}}(k)^2 + m_{\mathrm{II}}(k)^2 - 2m_{\mathrm{EE}}(k)m_{\mathrm{II}}(k) + 4m_{\mathrm{EI}}(k)m_{\mathrm{IE}}(k)} \quad . \tag{24}$$

The corresponding eigenvectors are

$$\boldsymbol{v}_{1,2}(k) = \mathcal{N}_{\pm} \begin{pmatrix} \sqrt{q}\, m_{\mathrm{EI}}(k) \\ m_{\pm}(k) - m_{\mathrm{EE}}(k) \end{pmatrix} \quad , \tag{25}$$

with normalization $\mathcal{N}_{\pm}$. The eigenvectors written in the Fourier basis are given by

$$\boldsymbol{v}_{\pm}(k) = \mathcal{N}_{\pm} \left[ \sqrt{q}\, m_{\mathrm{EI}}(k) \boldsymbol{v}_k^{(\mathrm{E})} + (m_{\pm}(k) - m_{\mathrm{EE}}(k))\, \boldsymbol{v}_k^{(\mathrm{I})} \right] \quad , \tag{26}$$

and we can get the eigenvectors $\tilde{\boldsymbol{v}}_{\pm}(k)$ in the basis we started with by extending $\boldsymbol{v}_k^{(\mathrm{E})}$ and $\boldsymbol{v}_k^{(\mathrm{I})}$ to vectors similar to Equation (22), where the elements corresponding to excitatory neurons are repeated $q$-times. Note that the normalization of the original basis leads to an additional factor $1/\sqrt{q}$ in the first term of Equation (26).

Analogously, we can find the left eigenvectors of $\boldsymbol{M}$ by conducting the same steps with the transpose of $\boldsymbol{M}$

$$\boldsymbol{u}_{\pm}(k) = \mathcal{N}_{\pm} \left[ m_{\mathrm{IE}}(k) \boldsymbol{v}_k^{(\mathrm{E})\dagger}/\sqrt{q} + (m_{\pm}(k) - m_{\mathrm{EE}}(k))\, \boldsymbol{v}_k^{(\mathrm{I})\dagger} \right] \quad , \tag{27}$$

and the vectors in the original basis $\tilde{\boldsymbol{u}}_{\pm}(k)$ are obtained similarly to the right eigenvectors. The normalization $\mathcal{N}_{\pm}$ is chosen such that

$$\begin{aligned}\tilde{\boldsymbol{u}}_{+}(k)\cdot\tilde{\boldsymbol{v}}_{+}(k)&=1 \quad ,\\ \tilde{\boldsymbol{u}}_{+}(k)\cdot\tilde{\boldsymbol{v}}_{-}(k)&=0 \quad ,\\ \tilde{\boldsymbol{u}}_{-}(k)\cdot\tilde{\boldsymbol{v}}_{+}(k)&=0 \quad ,\\ \tilde{\boldsymbol{u}}_{-}(k)\cdot\tilde{\boldsymbol{v}}_{-}(k)&=1 \quad ,\end{aligned}$$

which leads to

$$\mathcal{N}_{\pm}=\sqrt{m_{\mathrm{EI}}(k)m_{\mathrm{IE}}(k)-\left(m_{\pm}(k)-m_{\mathrm{EE}}(k)\right)^{2}} \quad .$$

Now, we can express $A^{-1}$ in terms of the eigenvalues and eigenvectors of $\boldsymbol{M}$

$$\boldsymbol{A}^{-1}=1+\sum_{k}\left(\frac{m_{+}(k)}{1-m_{+}(k)}\tilde{\boldsymbol{v}}_{+}(k)\cdot\tilde{\boldsymbol{u}}_{+}(k)+\frac{m_{-}(k)}{1-m_{-}(k)}\tilde{\boldsymbol{v}}_{-}(k)\cdot\tilde{\boldsymbol{u}}_{-}(k)\right) \quad , \tag{28}$$

which leads to

$$A_{ij}^{-1}=\delta_{ij}+\frac{1}{N}\sum_{k}\mu_{ij}(k)e^{\mathrm{i}k|\boldsymbol{x}_i-\boldsymbol{x}_j|} \quad , \tag{29}$$

where we defined $\boldsymbol{\mu}(k)$ similar to Equation (16). Let $E$ and $I$ be the sets of indices referring to excitatory or inhibitory neurons respectively. We find

$$\mu_{ij}(k)\equiv\begin{cases}\mu_{\mathrm{EE}}(k) & \text{for} \quad i,j\in E\\ \mu_{\mathrm{EI}}(k) & \text{for} \quad i\in E, j\in I\\ \mu_{\mathrm{IE}}(k) & \text{for} \quad i\in E, j\in I\\ \mu_{\mathrm{II}}(k) & \text{for} \quad i,j\in I\end{cases} \quad ,$$

with

$$\begin{aligned}\mu_{\mathrm{EE}}(k)&=\frac{1}{q}\frac{m_{\mathrm{EE}}(k)+m_{\mathrm{IE}}(k)m_{\mathrm{EI}}(k)-m_{\mathrm{EE}}(k)m_{\mathrm{II}}(k)}{1-\zeta(k)} \quad ,\\ \mu_{\mathrm{EI}}(k)&=\frac{m_{\mathrm{EI}}(k)}{1-\zeta(k)} \quad ,\\ \mu_{\mathrm{IE}}(k)&=\frac{1}{q}\frac{m_{\mathrm{IE}}(k)}{1-\zeta(k)} \quad ,\\ \mu_{\mathrm{II}}(k)&=\frac{m_{\mathrm{II}}(k)+m_{\mathrm{IE}}(k)m_{\mathrm{EI}}(k)-m_{\mathrm{EE}}(k)m_{\mathrm{II}}(k)}{1-\zeta(k)} \quad ,\end{aligned} \tag{30}$$

and

$$\zeta(k)=m_{\mathrm{EE}}(k)+m_{\mathrm{II}}(k)+m_{\mathrm{EI}}(k)m_{\mathrm{IE}}(k)-m_{\mathrm{EE}}(k)m_{\mathrm{II}}(k) \quad .$$

#### 9.2 General Results

The renormalized noise is evaluated using the same trick as in the one population case. We express the all-ones vector using eigenvectors of the variance matrix $\boldsymbol{S}$

$$I=a\tilde{\boldsymbol{v}}_{+}(0)+b\tilde{\boldsymbol{v}}_{-}(0) \quad .$$

Evaluating the coefficients $a$ and $b$ and inserting the corresponding solutions into Equation (10) yields

$$\boldsymbol{D}_{\mathrm{r}} = \mathrm{diag}\left(\underbrace{\underbrace{D_{\mathrm{r}}^{(\mathrm{E})},\ldots,D_{\mathrm{r}}^{(\mathrm{E})}}_{q-\text{times}},D_{\mathrm{r}}^{(\mathrm{I})},\underbrace{D_{\mathrm{r}}^{(\mathrm{E})},\ldots,D_{\mathrm{r}}^{(\mathrm{E})}}_{q-\text{times}},D_{\mathrm{r}}^{(\mathrm{I})},\ldots,\underbrace{D_{\mathrm{r}}^{(\mathrm{E})},\ldots,D_{\mathrm{r}}^{(\mathrm{E})}}_{q-\text{times}},D_{\mathrm{r}}^{(\mathrm{I})}}_{N(q+1)-\text{entries}}\right) \quad , \tag{31}$$

with

$$\begin{aligned} D_{\mathrm{r}}^{(\mathrm{E})} &= D\left[1+\frac{s_{\mathrm{EE}}(0)+s_{\mathrm{EI}}(0)+s_{\mathrm{EI}}(0)s_{\mathrm{IE}}(0)-s_{\mathrm{EE}}(0)s_{\mathrm{II}}(0)}{1-R^2}\right] \quad , \\ D_{\mathrm{r}}^{(\mathrm{I})} &= D\left[1+\frac{s_{\mathrm{IE}}(0)+s_{\mathrm{II}}(0)+s_{\mathrm{EI}}(0)s_{\mathrm{IE}}(0)-s_{\mathrm{EE}}(0)s_{\mathrm{II}}(0)}{1-R^2}\right] \quad , \end{aligned}$$

with the eigenvalues $s_{ab}(k)$ of $\boldsymbol{S}$. We again identified the spectral bound

$$R^2 = s_{\mathrm{EE}}(0)+s_{\mathrm{II}}(0)+s_{\mathrm{EI}}(0)s_{\mathrm{IE}}(0)-s_{\mathrm{EE}}(0)s_{\mathrm{II}}(0) \quad . \tag{32}$$

The mean covariances can be written as

$$\overline{\boldsymbol{c}} = \boldsymbol{D}_{\mathrm{r}} + \boldsymbol{\mu}\boldsymbol{D}_{\mathrm{r}} + \boldsymbol{D}_{\mathrm{r}}\boldsymbol{\mu}^{\mathrm{T}} + \boldsymbol{\mu}\boldsymbol{D}_{\mathrm{r}}\boldsymbol{\mu}^{\mathrm{T}} \quad ,$$

where $\boldsymbol{\mu} = \boldsymbol{\mu}(x)$. We can distinguish three different kinds of covariances depending on the type of neurons involved

$$\overline{c_{ij}} \equiv \begin{cases} \overline{c_{\mathrm{EE}}}(x) & \text{for} \quad i,j\in E \\ \overline{c_{\mathrm{EI}}}(x) & \text{for} \quad i\in E, j\in I \quad \text{or} \quad i\in E, j\in I \\ \overline{c_{\mathrm{II}}}(x) & \text{for} \quad i,j\in I \end{cases} \quad .$$

with

$$\begin{aligned} \overline{c_{\mathrm{EE}}}(x) &= D_{\mathrm{r}}^{(\mathrm{E})}\delta(x) + 2D_{\mathrm{r}}^{(\mathrm{E})}\mu_{\mathrm{EE}}(x) + D_{\mathrm{r}}^{(\mathrm{E})}q\left(\mu_{\mathrm{EE}}*\mu_{\mathrm{EE}}\right)(x) + D_{\mathrm{r}}^{(\mathrm{I})}\left(\mu_{\mathrm{EI}}*\mu_{\mathrm{EI}}\right)(x) \quad , \\ \overline{c_{\mathrm{EI}}}(x) &= D_{\mathrm{r}}^{(\mathrm{E})}\mu_{\mathrm{IE}}(x) + D_{\mathrm{r}}^{(\mathrm{I})}\mu_{\mathrm{EI}}(x) + D_{\mathrm{r}}^{(\mathrm{E})}q\left(\mu_{\mathrm{EE}}*\mu_{\mathrm{IE}}\right)(x) + D_{\mathrm{r}}^{(\mathrm{I})}\left(\mu_{\mathrm{II}}*\mu_{\mathrm{EI}}\right)(x) \quad , \\ \overline{c_{\mathrm{II}}}(x) &= D_{\mathrm{r}}^{(\mathrm{I})}\delta(x) + 2D_{\mathrm{r}}^{(\mathrm{I})}\mu_{\mathrm{II}}(x) + D_{\mathrm{r}}^{(\mathrm{E})}q\left(\mu_{\mathrm{IE}}*\mu_{\mathrm{IE}}\right)(x) + D_{\mathrm{r}}^{(\mathrm{I})}\left(\mu_{\mathrm{II}}*\mu_{\mathrm{II}}\right)(x) \quad . \end{aligned}$$

#### 9.3 Long-Range Limit

From here on, we consider the special case in which the synaptic connections only depend on the type of the presynaptic neuron and not on the type of the postsynaptic neuron. This is in agreement with network parameters used in established cortical network models (*69, 70*), in which the connection probabilities to both types of target neurons in the same layer are usually of the same order of magnitude. In that case, all expressions become independent of the first population index $A_{ab} \equiv A_b$, and the only expressions we need to evaluate become

$$\mu_a(k) = \gamma_a \frac{m_a(k)}{1-\zeta(k)} \quad ,$$

with

$$\zeta(k) = m_{\mathrm{E}}(k) + m_{\mathrm{I}}(k) \quad ,$$

and

$$\gamma_a = \begin{cases} 1 & \text{if } a = \mathrm{I} \\ 1/q & \text{if } a = \mathrm{E} \end{cases} \quad . \tag{33}$$

After taking the continuum limit, we can make a (0,2)-Padé approximation again

$$\mu_a(k) \approx \frac{\gamma_a m_a(0)}{1-\zeta(0) - \left[\frac{\zeta''(0)}{2} + (1-\zeta(0))\frac{m_a''(0)}{2m_a(0)}\right]k^2} \quad ,$$

which leads to the poles

$$k_0 = \pm\sqrt{\left[\frac{\zeta''(0)}{2\zeta(0)}\frac{\zeta(0)}{1-\zeta(0)} + \frac{m_a''(0)}{2m_a(0)}\right]^{-1}} \quad ,$$

or the effective decay constant of the mean covariances

$$\overline{d}_a = \mathrm{Im}(k_0)^{-1} = \sqrt{-\frac{\zeta''(0)}{2\zeta(0)}\frac{\zeta(0)}{1-\zeta(0)} - \frac{m_a''(0)}{2m_a(0)}} \quad .$$

Using

$$\begin{aligned} \zeta \equiv \zeta(0) &= w_\mathrm{E}K_\mathrm{E} + w_\mathrm{I}K_\mathrm{I} \quad , \\ \zeta'' \equiv \zeta''(0) &= -w_\mathrm{E}K_\mathrm{E}\left\langle x^2\right\rangle_\mathrm{E} - w_\mathrm{I}K_\mathrm{I}\left\langle x^2\right\rangle_\mathrm{I} \quad , \\ m_a(0) &= w_a K_a \quad , \\ m_a''(0) &= -w_a K_a \left\langle x^2\right\rangle_a \quad , \end{aligned}$$

we get

$$\begin{aligned} \overline{d}_a &= \sqrt{\frac{w_\mathrm{E}K_\mathrm{E}\left\langle x^2\right\rangle_\mathrm{E} + w_\mathrm{I}K_\mathrm{I}\left\langle x^2\right\rangle_\mathrm{I}}{w_\mathrm{E}K_\mathrm{E} + w_\mathrm{I}K_\mathrm{I}}\frac{\zeta}{1-\zeta} + \frac{\left\langle x^2\right\rangle_a}{2}} \\ &= \sqrt{\frac{(\omega\kappa\tilde{\eta}^2+1)}{\omega\kappa+1}\frac{\zeta}{1-\zeta}\frac{\left\langle x^2\right\rangle_\mathrm{I}}{2} + \frac{\left\langle x^2\right\rangle_a}{2}} \quad , \end{aligned}$$

after introducing relative parameters

$$\omega = \frac{w_\mathrm{E}}{w_\mathrm{I}}, \quad \kappa = \frac{K_\mathrm{E}}{K_\mathrm{I}}, \quad \tilde{\eta}^2 = \frac{\left\langle x^2\right\rangle_\mathrm{E}}{\left\langle x^2\right\rangle_\mathrm{I}}, \quad \eta = \frac{\lambda_\mathrm{E}}{\lambda_\mathrm{I}} \quad .$$

The renormalized noise Equation (10) reduces to

$$D_\mathrm{r} = \frac{D}{1-R^2} \quad . \tag{34}$$

The mean covariances are

$$\begin{aligned} \overline{c_\mathrm{EE}}(x) &= D_\mathrm{r}\left[\delta(x) + 2\mu_\mathrm{E}(x) + q\left(\mu_\mathrm{E} * \mu_\mathrm{E}\right)(x) + \left(\mu_\mathrm{I} * \mu_\mathrm{I}\right)(x)\right] \quad , \\ \overline{c_\mathrm{EI}}(x) &= D_\mathrm{r}\left[\mu_\mathrm{E}(x) + \mu_\mathrm{I}(x) + q\left(\mu_\mathrm{E} * \mu_\mathrm{E}\right)(x) + \left(\mu_\mathrm{I} * \mu_\mathrm{I}\right)(x)\right] \quad , \\ \overline{c_\mathrm{II}}(x) &= D_\mathrm{r}\left[\delta(x) + 2\mu_\mathrm{I}(x) + q\left(\mu_\mathrm{E} * \mu_\mathrm{E}\right)(x) + \left(\mu_\mathrm{I} * \mu_\mathrm{I}\right)(x)\right] \quad , \end{aligned}$$

with

$$\mu_a(x) = \gamma_a \frac{m_a(0)}{2(1-\zeta)\overline{d}_a} \exp\left(-\frac{|x|}{\overline{d}_a}\right) \quad ,$$

and

$$\left(\mu_a * \mu_a\right)(x) = \left(\gamma_a \frac{m(0)}{2\left(1-\zeta\right)\overline{d}_a}\right)^2 \left(\overline{d}_a + |x|\right) \exp\left(-\frac{|x|}{\overline{d}_a}\right) \quad .$$

Note that expressions coming from both populations contribute to each kind of covariance. Therefore, all mean covariances contain a part that decays with either of the decay constants we just determined. If, for example, the inhibitory decay constant is much larger than the excitatory one, $\overline{c_{\mathrm{EI}}}(x)$ will decay with the largest decay constant in the long-range limit

**Exponential Connectivity Profile:** Just as in Section 8.4 we get

$$\overline{d}_a = \sqrt{\frac{(\omega\kappa\eta^2+1)}{\omega\kappa+1}\frac{\lambda_0}{1-\lambda_0}d_{\mathrm{I}}^2 + d_a^2} \quad , d_{\mathrm{eff},a} = \sqrt{\frac{(\omega^2\kappa\eta^2+1)}{\omega^2\kappa+1}\frac{R^2}{1-R^2}d_{\mathrm{I}}^2 + d_a^2} \quad ,$$

with $\lambda_0 = w_{\mathrm{E}}K_{\mathrm{E}} + w_{\mathrm{I}}K_{\mathrm{I}}$ for the decay constant of the mean covariances, and $R^2 = w_{\mathrm{E}}^2 K_{\mathrm{E}} + w_{\mathrm{I}}^2 K_{\mathrm{I}}$ for the decay constant of the variances.

**Gaussian Connectivity Profile:** And similar to Section 8.4 we get

$$\overline{d}_a = \sqrt{\frac{(\omega\kappa\eta^2+1)}{\omega\kappa+1}\frac{\lambda_0}{1-\lambda_0}\frac{d_{\mathrm{I}}^2}{2} + \frac{d_a^2}{2}} \quad , d_{\mathrm{eff},a} = \sqrt{\frac{(\omega\kappa\eta^2+1)}{\omega\kappa+1}\frac{\lambda_0}{1-\lambda_0}\frac{d_{\mathrm{I}}^2}{2} + \frac{d_a^2}{2}} \quad .$$

### 10 Two-Dimensional Network with One Population

In the following, we are considering two-dimensional networks, which are supposed to mimic a single-layered cortical network. Neurons are positioned on a two-dimensional lattice ($N_x \times N_y$ grid) with periodic boundary conditions in both dimensions (a torus). We define the activity vector to be of the form

$$\boldsymbol{x} = \begin{pmatrix} x_{1,1} \\ x_{1,2} \\ \vdots \\ x_{1,N_y} \\ x_{2,1} \\ \vdots \\ x_{2,N_y} \\ \vdots \\ x_{N_x,1} \\ \vdots \\ x_{N_x,N_y} \end{pmatrix} \quad .$$

The connectivity matrix is defined correspondingly.

#### 10.1 Dimensionality Reduction

In two dimensions we have to define two translation operators that move all neurons either one step in the $x$-direction, or the $y$-direction, respectively. They are defined via their action on $\boldsymbol{x}$

$$\boldsymbol{T}_x\boldsymbol{x} = \begin{pmatrix} x_{N_x,1} \\ x_{N_x,2} \\ \vdots \\ x_{N_x,N_y} \\ x_{1,1} \\ \vdots \\ x_{1,N_y} \\ \vdots \\ x_{N_x-1,1} \\ \vdots \\ x_{N_x-1,N_y} \end{pmatrix} \quad , \boldsymbol{T}_y\boldsymbol{x} = \begin{pmatrix} x_{1,N_y} \\ x_{1,1} \\ \vdots \\ x_{1,N_y-1} \\ x_{2,N_y} \\ \vdots \\ x_{2,N_y-1} \\ \vdots \\ x_{N_x,N_y} \\ \vdots \\ x_{N_x,N_y-1} \end{pmatrix} \quad . \tag{35}$$

Similar reasoning as in one dimension leads to the eigenvalues

$$\mathrm{e}^{-\mathrm{i}k_l^{(x)}a} \quad , \quad k_l^{(x)} = \frac{2\pi}{L_x}l \quad , \quad l \in \{0, 1, ..., N_x - 1\} \quad ,$$

and similar for the $y$-direction. The eigenvectors can be inferred from the recursion relations

$$\begin{aligned} \boldsymbol{T}_x\boldsymbol{v} &= \mathrm{e}^{-\mathrm{i}k_l^{(x)}a}\boldsymbol{v} \quad \Rightarrow \quad v_{(\alpha+1)\beta} = \mathrm{e}^{\mathrm{i}k_l^{(x)}a}v_{\alpha\beta} \quad , \\ \boldsymbol{T}_y\boldsymbol{v} &= \mathrm{e}^{-\mathrm{i}k_l^{(y)}a}\boldsymbol{v} \quad \Rightarrow \quad v_{\alpha(\beta+1)} = \mathrm{e}^{\mathrm{i}k_l^{(y)}a}v_{\alpha\beta} \quad , \end{aligned}$$

where entries $v_{\alpha\beta}$ of the vector $\boldsymbol{v}$ are defined analogously to Equation (35). The eigenvectors are given by

$$\boldsymbol{v}_{\boldsymbol{k}} = \frac{1}{\sqrt{N_xN_y}} \begin{pmatrix} \boldsymbol{v}^{(x)} \\ \mathrm{e}^{\mathrm{i}k^{(y)}a}\boldsymbol{v}^{(x)} \\ \vdots \\ \mathrm{e}^{-\mathrm{i}k^{(y)}a}\boldsymbol{v}^{(x)} \end{pmatrix} \quad , \quad \boldsymbol{v}^{(x)} = \begin{pmatrix} 1 \\ \mathrm{e}^{\mathrm{i}k^{(x)}a} \\ \vdots \\ \mathrm{e}^{\mathrm{i}\frac{N_x-1}{2}k^{(x)}a} \\ \mathrm{e}^{-\mathrm{i}\frac{N_x-1}{2}k^{(x)}a} \\ \vdots \\ \mathrm{e}^{-\mathrm{i}k^{(x)}a} \end{pmatrix} \quad ,$$

where we suppressed the subscripts of $k^{(x)}$ and $k^{(y)}$ again. Using that these eigenvectors are eigenvectors of $\boldsymbol{M}$ as well, yields the eigenvalues of $\boldsymbol{M}$

$$m_{\boldsymbol{k}} = \boldsymbol{v}_{\boldsymbol{k}}^{\dagger}\boldsymbol{M}\boldsymbol{v}_{\boldsymbol{k}} = Kw\sum_x\sum_y p(|\boldsymbol{x}|)\mathrm{e}^{-\mathrm{i}\boldsymbol{k}\cdot\boldsymbol{x}} \quad .$$

In the continuum limit, this becomes the two-dimensional Fourier transform

$$m(\boldsymbol{k}) = Kw\int \mathrm{d}^2x\, p(\boldsymbol{x})\mathrm{e}^{-\mathrm{i}\boldsymbol{k}\cdot\boldsymbol{x}} \quad . \tag{36}$$

The inverse of $A$ is given by

$$A^{-1}(\boldsymbol{x}) = \delta(\boldsymbol{x}) + \mu(\boldsymbol{x}) \quad , \tag{37}$$

with the inverse two-dimensional Fourier transform

$$\mu(\boldsymbol{x}) = \frac{1}{(2\pi)^2} \int \mathrm{d}^2 k \, \frac{m(\boldsymbol{k})}{1 - m(\boldsymbol{k})} \mathrm{e}^{\mathrm{i}\boldsymbol{k}\cdot\boldsymbol{x}} \quad . \tag{38}$$

The expression for the renormalized noise is the same as in the one-dimensional case with one population. Hence, the mean covariances are given by

$$\bar{c}(\boldsymbol{x}) = \frac{D}{1 - R^2} \left[\delta(\boldsymbol{x}) + 2\mu(\boldsymbol{x}) + (\mu * * \mu)(\boldsymbol{x})\right] \quad , \tag{39}$$

which is the one-dimensional expression, except for the convolution, which is replaced by its two-dimensional analogon denoted here by the double asterisk.

#### 10.2 Long-Range Limit

Employing the symmetry of the connectivity kernel, we rewrite the integral in $\mu(\boldsymbol{x})$ using polar coordinates

$$\mu(\boldsymbol{x}) = \frac{1}{(2\pi)^2} \int_0^\infty \mathrm{d}k \int_0^{2\pi} \mathrm{d}\varphi \, k \frac{m(k)}{1 - m(k)} \mathrm{e}^{\mathrm{i}kr\cos(\varphi)} \quad , \tag{40}$$

with $r = |\boldsymbol{x}|$, and make a Padé approximation of order (0,2) of the integration kernel

$$\mu(\boldsymbol{x}) = \frac{1}{(2\pi)^2} \int_0^\infty \mathrm{d}k \int_0^{2\pi} d\varphi \, k \frac{m(0)}{1 - m(0) - \frac{m''(0)}{2m(0)} k^2} \mathrm{e}^{\mathrm{i}kr\cos(\varphi)} \quad . \tag{41}$$

Following (*71, p.160f*), we can interpret this as calculating the Green's function of the heat equation

$$\left[1 - m(0) + \frac{m''(0)}{2m(0)} \nabla^2\right] \mu(\boldsymbol{x}) = m(0)\delta(r) \quad , \tag{42}$$

which can be solved, using the fact that $\mu(\boldsymbol{x})$ can only be a function of the radial distance $r$, due to the given symmetry of the kernel. Rewriting leads to

$$\left[-\frac{1}{r}\frac{\mathrm{d}}{\mathrm{d}r}\left(r\frac{\mathrm{d}}{\mathrm{d}r}\right) + \bar{d}^{-2}\right] \mu(r) = \Gamma\delta(r) \quad ,$$

with the effective decay constant

$$\bar{d} = \sqrt{-\frac{m''(0)}{2m(0)} \frac{1}{1 - m(0)}} \quad , \tag{43}$$

and $\Gamma = -2m(0)^2/m''(0)$. Defining $\rho \equiv r/\bar{d}$, $\tilde{\mu}(\rho) \equiv \mu(r/\bar{d})$, and using $\delta(\rho\bar{d}) = \bar{d}^{-2}\delta(\rho)$, we get

$$\left[-\frac{1}{\rho}\frac{\mathrm{d}}{\mathrm{d}\rho}\left(\rho\frac{\mathrm{d}}{\mathrm{d}\rho}\right) + 1\right] \tilde{\mu}(\rho) = \Gamma\delta(\rho) \quad .$$

The solution to this equation is given by the modified Bessel function of second kind and zeroth order $K_0$

$$\tilde{\mu}(\rho) = \frac{\Gamma}{2\pi} K_0(\rho) \quad .$$

Reinserting the defined variables yields

$$\mu(r) = -\frac{m(0)^2}{\pi m''(0)} K_0\left(\frac{r}{\bar{d}}\right) \quad . \tag{44}$$

Note that the modified Bessel functions of second kind decay exponentially for long distances

$$K_i\left(\frac{r}{\bar{d}}\right) \xrightarrow{r\to\infty} \sqrt{\frac{\pi \bar{d}}{2r}}\mathrm{e}^{-r/\bar{d}} \quad . \tag{45}$$

But, consider that the inverse square root of the distance appears in front of the exponential. Formally, this is the one-dimensional result. The only difference here is, that $m(k)$ is a two-dimensional Fourier transform instead of a one-dimensional one and $\mu(r)$ contains modified Bessel functions of second kind instead of exponentials.

In order to evaluate the expression for the mean covariances (39), one needs to calculate the two-dimensional convolution of a modified Bessel function of second kind with itself, for which we use the following trick

$$\begin{aligned}
(K_0 ** K_0)\left(\frac{r}{\bar{d}}\right) &= \mathcal{F}^{-1}\left[\tilde{K}_0 \cdot \tilde{K}_0\right]\left(\frac{r}{\bar{d}}\right) \\
&= \frac{1}{2\pi}\mathcal{H}^{-1}\left[\frac{1}{(\beta + k^2)^2}\right]\left(\sqrt{\beta} r\right) \\
&= -\frac{1}{2\pi}\frac{\mathrm{d}}{\mathrm{d}\beta}\mathcal{H}^{-1}\left[\frac{1}{\beta + k^2}\right]\left(\sqrt{\beta} r\right) \\
&= -\frac{1}{2\pi}\frac{\mathrm{d}}{\mathrm{d}\beta}K_o\left(\sqrt{\beta} r\right) \\
&= \frac{d_{\mathrm{eff},\mu} r}{4\pi}K_1\left(\frac{r}{\bar{d}}\right) \quad ,
\end{aligned}$$

where $\mathcal{F}$ denotes the Fourier transform, $\mathcal{H}$ denotes the Hankel transform, and $\beta = \bar{d}^{-2}$. The last step can be found in (*72, 9.6.27*).

The mean covariances are given by

$$\begin{aligned}
\bar{c}(r) \quad &= \quad \frac{D}{1-R^2}\left[\delta(r) - 2\frac{m(0)^2}{\pi m''(0)}K_0\left(\frac{r}{\bar{d}}\right) + \frac{m(0)^4}{m''(0)^2}\frac{\bar{d} r}{4\pi^3}K_1\left(\frac{r}{\bar{d}}\right)\right] \\
&\xrightarrow{r\to\infty} \quad \frac{D}{1-R^2}\left[\delta(r) - \frac{m(0)^2}{m''(0)}\sqrt{\frac{2\bar{d}}{\pi r}}\mathrm{e}^{-r/\bar{d}} + \frac{m(0)^4}{m''(0)^2}\sqrt{\frac{\bar{d}^3 r}{32\pi^5}}\mathrm{e}^{-r/\bar{d}}\right] \quad .
\end{aligned}$$

Using

$$m(0) = Kw \equiv \zeta \quad , \quad m''(0) = -Kw\left\langle r^2\right\rangle \quad , \tag{46}$$

we get the effective decay constant

$$\bar{d} = \sqrt{\frac{1}{1-\zeta}\frac{\langle r^2\rangle}{2}}d \quad . \tag{47}$$

**Exponential Connectivity Profile:** Using a two-dimensional exponential connectivity profile

$$p(\boldsymbol{x}) = \frac{1}{2\pi d^2}\mathrm{e}^{-|\boldsymbol{x}|/d} \quad ,$$

leads to $\left\langle r^2 \right\rangle = 6d^2$, and we get

$$\bar{d} = \sqrt{\frac{3}{1-\lambda_0}}d \quad , d_{\text{eff}} = \sqrt{\frac{3}{1-R^2}}d \quad ,$$

with $\lambda_0 = Kw$, and $R^2 = Kw^2$.

**Gaussian Connectivity Profile:** Using a two-dimensional Gaussian connectivity profile

$$p(\boldsymbol{x}) = \frac{1}{2\pi d^2}\mathrm{e}^{-\boldsymbol{x}^2/(2d^2)} \quad ,$$

leads to $\left\langle r^2 \right\rangle = 2d^2$, and we get

$$\bar{d} = \sqrt{\frac{1}{1-\lambda_0}}d \quad , d_{\text{eff}} = \sqrt{\frac{1}{1-R^2}}d \quad .$$

#### 10.3 Note on Higher Order Approximation

While the (0,2)-Padé approximation seems to yield good results for the one-dimensional cases, in two dimensions the results only coincide for large spectral radii (Appendix 1-figure 4). One can extract a higher order approximation of the poles of the integration kernel of $\mu(\boldsymbol{x})$ and thereby the effective decay constant $d_{\text{eff}}$ using the DLog-Pade-method, for which one calculates an $(n, n+1)$-Padé approximation of the logarithmic derivative of the integration kernel around zero (*73*). Using a (1,2)Padé approximation leads to

$$\bar{d} = \sqrt{-\frac{3(2m(0)-1)m''(0)^2 + (1-m(0))m(0)m''''(0)}{6m''(0)m(0)(1-m(0))}} \quad ,$$

which coincides with our previous results in the limit $m(0) \to 1$, and thus for large spectral radii. Note that this expression contains the fourth moment of the connectivity kernel $m''''(0) = wK\langle x^4 \rangle$.

### 11 Two-Dimensional Network with Two Populations

Finally, we consider a two-dimensional network with two populations of neurons. As in the one dimensional case, the neurons are gathered in cells, which contain one inhibitory and $q$ excitatory neurons. Again, they are placed on a two-dimensional lattice with periodic boundary conditions. The activity vector takes the form

$$\boldsymbol{x} = \begin{pmatrix} \boldsymbol{x}_{1,1}^{(\mathrm{E})} \\ x_{1,1}^{(\mathrm{I})} \\ \boldsymbol{x}_{1,2}^{(\mathrm{E})} \\ x_{1,2}^{(\mathrm{I})} \\ \vdots \\ \boldsymbol{x}_{1,N_y}^{(\mathrm{E})} \\ x_{1,N_y}^{(\mathrm{I})} \\ \boldsymbol{x}_{2,1}^{(\mathrm{E})} \\ x_{2,1}^{(\mathrm{I})} \\ \vdots \\ \boldsymbol{x}_{N_x,N_y}^{(\mathrm{E})} \\ x_{N_x,N_y}^{(\mathrm{I})} \end{pmatrix} \quad , \tag{48}$$

where $\boldsymbol{x}_{i,j}^{(\mathrm{E})}$ denotes a $q$-dimensional vector.

##### 11.1 Dimensionality Reduction

We apply the procedure developed so far, which leads to the results we found in the one-dimensional case with two populations, with Fourier transforms and convolutions replaced by their two-dimensional analogons and modified Bessel functions of second kind instead of exponentials. So, we end up with

$$\begin{aligned}\overline{c_{\mathrm{EE}}}(x) &= D_{\mathrm{r}}^{(\mathrm{E})}\delta(x) + 2D_{\mathrm{r}}^{(\mathrm{E})}\mu_{\mathrm{EE}}(x) + D_{\mathrm{r}}^{(\mathrm{E})}q\left(\mu_{\mathrm{EE}} ** \mu_{\mathrm{EE}}\right)(x) + D_{\mathrm{r}}^{(\mathrm{I})}\left(\mu_{\mathrm{EI}} ** \mu_{\mathrm{EI}}\right)(x) \quad ,\\ \overline{c_{\mathrm{EI}}}(x) &= D_{\mathrm{r}}^{(\mathrm{E})}\mu_{\mathrm{IE}}(x) + D_{\mathrm{r}}^{(\mathrm{I})}\mu_{\mathrm{EI}}(x) + D_{\mathrm{r}}^{(\mathrm{E})}q\left(\mu_{\mathrm{EE}} ** \mu_{\mathrm{IE}}\right)(x) + D_{\mathrm{r}}^{(\mathrm{I})}\left(\mu_{\mathrm{II}} ** \mu_{\mathrm{EI}}\right)(x) \quad ,\\ \overline{c_{\mathrm{II}}}(x) &= D_{\mathrm{r}}^{(\mathrm{I})}\delta(x) + 2D_{\mathrm{r}}^{(\mathrm{I})}\mu_{\mathrm{II}}(x) + D_{\mathrm{r}}^{(\mathrm{E})}q\left(\mu_{\mathrm{IE}} ** \mu_{\mathrm{IE}}\right)(x) + D_{\mathrm{r}}^{(\mathrm{I})}\left(\mu_{\mathrm{II}} ** \mu_{\mathrm{II}}\right)(x) \quad ,\end{aligned}$$

and $\mu_{ab}(x)$ given by (30) and the two-dimensional Fourier transform

$$m_{ab}(\boldsymbol{k}) = K_{ab}w_{ab}\int \mathrm{d}^2x\, p_{ab}(\boldsymbol{x})\mathrm{e}^{-\mathrm{i}\boldsymbol{k}\cdot\boldsymbol{x}} \quad .$$

The renormalized noise is given by (31) with spectral bound (32), with the eigenvalues $s_{ab}(k)$ replaced by the two-dimensional Fourier transforms $s_{ab}(\boldsymbol{k})$.

##### 11.2 Long-Range Limit

Again, considering the special case in which the synaptic connections only depend on the type of the presynaptic neuron and not on the type of the postsynaptic neuron, the expressions simplify to

$$\mu_a(k) = \frac{m_a(k)}{1-\zeta(k)} \quad , \tag{49}$$

with

$$\zeta(k) = m_{\mathrm{E}}(k) + m_{\mathrm{I}}(k) \quad .$$

Padé approximation of the Fourier kernel, integration using (*71, p.160f*) and suppressing the zero arguments of $\zeta$ and $m_a$ leads to

$$\begin{aligned}\mu_a(r) \quad &= \quad -\frac{\gamma_a w_a K_a}{2\pi\left(1-\zeta\right)\bar{d}_a^2}\mathrm{K}_0\left(\frac{r}{\bar{d}_a}\right) \quad , \\ &\xrightarrow{r\to\infty} \quad -\frac{\gamma_a w_a K_a}{(1-\zeta)}\sqrt{\frac{1}{8\pi r d_{\mathrm{eff},\mu,a}^3}}\mathrm{e}^{-r/d_{\mathrm{eff},\mu,\mathrm{a}}}\end{aligned} \tag{50}$$

with

$$\bar{d}_a = \sqrt{-\frac{\zeta''}{2\zeta}\frac{\zeta}{1-\zeta} - \frac{m_a''}{2m_a}} \quad .$$

After introducing the same relative parameters as in Section 9.3, we find

$$\bar{d}_a = \sqrt{\frac{(\omega\kappa\tilde{\eta}^2+1)}{\omega\kappa+1}\frac{\zeta}{1-\zeta}\frac{\langle x^2\rangle_{\mathrm{I}}}{2} + \frac{\langle x^2\rangle_a}{2}} \quad . \tag{51}$$

The two-dimensional convolutions are given by

$$\begin{aligned}\left(\mu_a * * \mu_a\right)(r) &= \left[\frac{\gamma_a w_a K_a}{4(1-\zeta)}\right]^2 \frac{1}{\pi^3 \overline{d}_a^3} K_1\left(\frac{r}{\overline{d}_a}\right) \\ &\xrightarrow{r\to\infty} \left[\frac{\gamma_a w_a K_a}{4(1-\zeta)}\right]^2 \sqrt{\frac{1}{2\pi^5 \overline{d}_a^5 r}}\, \mathrm{e}^{-r/\overline{d}_a} \quad .\end{aligned} \tag{52}$$

The renormalized noise simplifies to (34). The mean covariances are given by

$$\begin{aligned}\overline{c_{\mathrm{EE}}}(x) &= D_{\mathrm{r}}\left[\delta(x) + 2\mu_{\mathrm{E}}(x) + q\left(\mu_{\mathrm{E}} * * \mu_{\mathrm{E}}\right)(x) + \left(\mu_{\mathrm{I}} * * \mu_{\mathrm{I}}\right)(x)\right] \quad , \\ \overline{c_{\mathrm{EI}}}(x) &= D_{\mathrm{r}}\left[\mu_{\mathrm{E}}(x) + \mu_{\mathrm{I}}(x) + q\left(\mu_{\mathrm{E}} * * \mu_{\mathrm{E}}\right)(x) + \left(\mu_{\mathrm{I}} * * \mu_{\mathrm{I}}\right)(x)\right] \quad , \\ \overline{c_{\mathrm{II}}}(x) &= D_{\mathrm{r}}\left[\delta(x) + 2\mu_{\mathrm{I}}(x) + q\left(\mu_{\mathrm{E}} * * \mu_{\mathrm{E}}\right)(x) + \left(\mu_{\mathrm{I}} * * \mu_{\mathrm{I}}\right)(x)\right] \quad .\end{aligned} \tag{53}$$

Remember that the result for the variances of the covariances is obtained by substituting $D_{\mathrm{r}}$ by its square, and $w_a$, or $\omega$ respectively, by its square and setting $\zeta = R^2$.

Equation (2) in the main text can be proven by inserting the result for $\overline{d}_a$

$$\overline{d}_{\mathrm{E}}^2 - \overline{d}_{\mathrm{I}}^2 = \frac{\left\langle x^2\right\rangle_{\mathrm{E}}}{2} - \frac{\left\langle x^2\right\rangle_{\mathrm{I}}}{2} = \mathrm{const.} \cdot \left(d_{\mathrm{E}}^2 - d_{\mathrm{I}}^2\right) \quad .$$

Using an exponential connectivity profile yields $\mathrm{const.} = 3$, a Gaussian connectivity profile yields $\mathrm{const.} = 1$.

**Exponential Connectivity Profile:** Using the results from 10.2, we find

$$\overline{d}_a = \sqrt{3\left[\frac{(\omega\kappa\eta^2+1)}{\omega\kappa+1}\frac{\lambda_0}{1-\lambda_0}d_I + d_a\right]} \quad , d_{\mathrm{eff},a} = \sqrt{3\left[\frac{(\omega^2\kappa\eta^2+1)}{\omega^2\kappa+1}\frac{R^2}{1-R^2}d_I + d_a\right]} \quad ,$$

with $\lambda_0 = w_{\mathrm{E}}K_{\mathrm{E}} + w_{\mathrm{I}}K_{\mathrm{I}}$, and $R^2 = w_{\mathrm{E}}^2 K_{\mathrm{E}} + w_{\mathrm{I}}^2 K_{\mathrm{I}}$.

**Gaussian Connectivity Profile:** Using the results from 10.2, we find

$$\overline{d}_a = \sqrt{\frac{(\omega\kappa\eta^2+1)}{\omega\kappa+1}\frac{\lambda_0}{1-\lambda_0}d_I + d_a} \quad , d_{\mathrm{eff},a} = \sqrt{\frac{(\omega^2\kappa\eta^2+1)}{\omega^2\kappa+1}\frac{R^2}{1-R^2}d_I + d_a} \quad .$$

#### 11.3 Higher Order Approximation

Using a (1,2)-DLog-Padé method as in Section 10.3 yields

$$\overline{d}_a = \sqrt{-\frac{(1-\zeta)^2\left(m_a m_a'''' - 3m_a''^2\right) + m_a^2\left[(1-\zeta)\zeta'''' + 3\zeta''^2\right]}{6m_a(1-\zeta)\left[(1-\zeta)m_a'' + m\zeta''\right]}} \quad , \tag{54}$$

which again contains the fourth moments of the connectivity kernels.

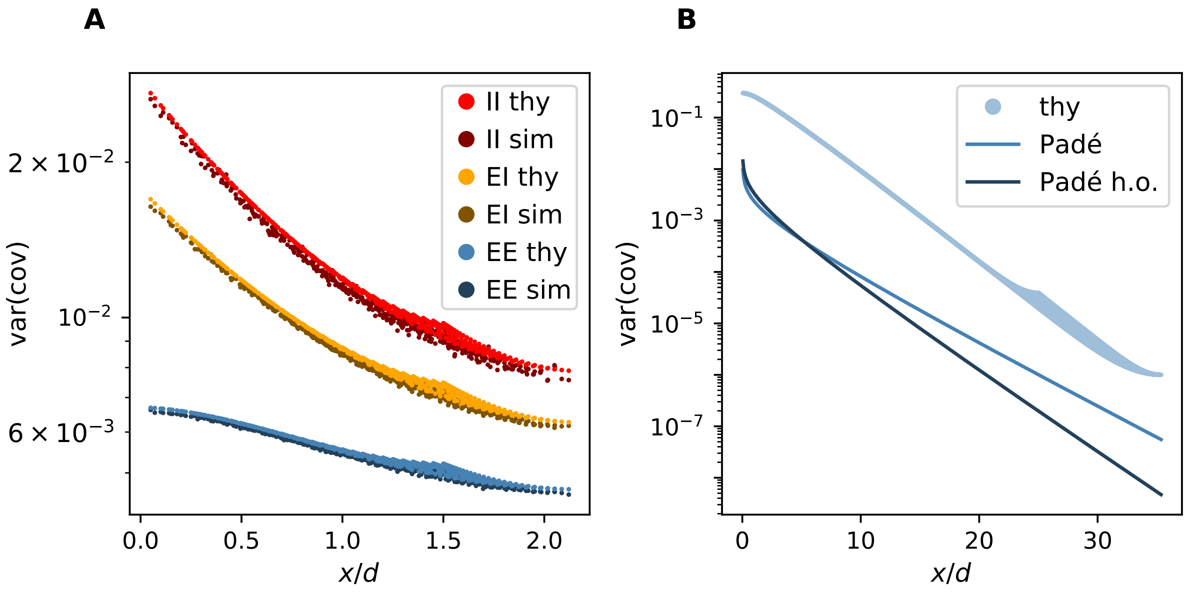


Appendix 1-figure 4: **Comparison of simulation and theory.** (**a**) Variance of EE, EI, and II covariances as a function of distance. Darker dots are the results of the simulation. Lighter ones are the prediction of the discrete theory. (**b**) Variance of EE covariances as a function of distance (Equation (53) for variances). The lightest blue dots are the predictions of the discrete theory ($\mu_a$ replaced by the discrete Fourier transform of Equation (49), taking into account Section 6), the medium blue line is the (0,2)-Padé prediction ($\mu_a$ replaced by its Padé approximation Equation (50), taking into account Section 6), and the dark blue line is the higher order (1,2)-DLog-Padé prediction ($\mu_a$ replaced by its Padé approximation Equation (50), using Equation (54), and taking into account Section 6). The source data and code used to create this figure are provided as supp_fig4.zip.

### 12 Validation of Theory

In order to validate our results, we performed simulations, in which an effective connectivity matrix $\boldsymbol{W}$ of a two-dimensional network was drawn randomly, and covariances were calculated using the result from (*35, 24, 31*)

$$\boldsymbol{c}(\boldsymbol{W}) = (1-\boldsymbol{W})^{-1}\,\boldsymbol{D}\,(1-\boldsymbol{W})^{-\mathrm{T}} \quad .$$

The elements of the different components $\boldsymbol{W}_{ab}$ of the effective connectivity matrix, similar to Equation (21), were drawn from a binomial distribution with $K_b$ trials and a success probability of $\gamma_b p_b(|\boldsymbol{x}|)$, with $\gamma_b$ given by Equation (33) and $|\boldsymbol{x}|$ denoting the distance between the neurons.

We compared the results to the predictions by our discrete theory, continuum theory, and the long-range limit. We did this for all cases presented above: one dimension with one population, one dimension with two populations, two dimensions with one population, and two dimensions with two populations. In the cases of two populations we solely considered the special case of synaptic connections only depending on the type of the presynaptic neuron. The first three cases are not reported here. We simulated several sets of parameters, varying the number of neurons, the number of inputs, the decay constants and the spectral bound, of which we only report the one using the parameters listed in Appendix 1-table 3, because the results do not differ qualitatively. Using

$$R^2 = s(0) = K_\mathrm{E} w_\mathrm{E}^2 + K_\mathrm{I} w_\mathrm{I}^2 \quad ,$$

| | Fig. 3 *B*,*C* | Fig. 3 *D*,*E* | Fig. 4 *A* | Fig. 4 *B* | |
|---|---|---|---|---|---|
| $N_x$ | 61 | 201 | 61 | 1001 | Number of neurons in x-direction |
| $N_y$ | 61 | 201 | 61 | 1001 | Number of neurons in y-direction |
| $q$ | 4 | 4 | 4 | 4 | Ratio of excitatory to inhibitory neurons |
| $K_{\mathrm{E}}$ | 100 | 100 | 100 | 100 | Number of excitatory inputs per neuron |
| $K_{\mathrm{I}}$ | 50 | 50 | 50 | 50 | Number of inhibitory inputs per neuron |
| $d_{\mathrm{E}}$ | 20 | 20 | 20 | 20 | Decay constant of excitatory connectivity profile |
| $d_{\mathrm{I}}$ | 10 | 10 | 10 | 10 | Decay constant of inhibitory connectivity profile |
| $D$ | 1 | 1 | 1 | 1 | Squared noise amplitude |
| $R$ | 0.95 | 0.95 | 0.8 | 0.95 | Spectral bound |
| | exponential | exponential | exponential | exponential | Connectivity kernel |

Appendix 1-table 3: Parameters used to create theory figures. Decay constants in units of lattice constant $a$.

and choosing

$$\frac{w_{\mathrm{I}}}{w_{\mathrm{E}}} = -\frac{N_E}{N_I} = -q \quad ,$$

we calculated the synaptic weights

$$w_{\mathrm{E}} = \frac{R}{\sqrt{K_{\mathrm{E}} + q^2 K_{\mathrm{I}}}} \quad , \quad w_{\mathrm{I}} = -\frac{qR}{\sqrt{K_{\mathrm{E}} + q^2 K_{\mathrm{I}}}} \quad .$$

The comparison of simulation and discrete theory is shown in Appendix 1-figure 4a. Simulation and theory match almost perfectly. The continuum theory, which is shown in figure 3*D*,*E* of the main text, matches as well as the discrete theory (not shown here). The slight shift in y-direction in Appendix 1-figure 4a is due to the fact that in the random realization of the network the spectral bound is not exactly matching the desired value, but is slightly different for each realization and distributed around the chosen value. This jittering around the real spectral bound is more pronounced as $R \to 1$. Note that the simulated networks were small compared to the decay constant of the connectivity profile, in order to keep simulation times reasonable. This is why the variances do not fall off linearly in the semi-log plot. The spreading at the end of the curves is a finite size effect due to periodic boundary conditions. The kink around $x/d = 1.25$ coincides with the size of the network in both $x$-, and $y$-direction.

In order to validate the long-range limit, we compared our discrete theory with the result from the Padé approximation at large distances (Appendix 1-figure 4b). We do not expect the Padé approximation to hold at small distances. The slope at large distances for the (0,2)-Padé approximation is smaller than the prediction by our theory, but the higher order approximation matches our theory very well. In the limit $R \to 1$ both Padé predictions yield similar results. The absolute value of the Padé approximation can be obtained from a residue analysis. This leads to values which are slightly off. The mismatch is no problem for the analysis here, which only focuses on the decay constant. Note that for plotting the higher order prediction in Appendix 1-figure 4b, we just inserted Equation (54) into Equation (50) and Equation (52). Calculating the correct residues for the (1,2)-DLog Padé approximation would lead to a better approximation of the absolute value.

### 13 NEST Simulation

For illustrating the dynamic nature of covariances emerging in the dynamically balanced critical state, we performed a simulation with the neural simulation tool NEST (*74*) using the parameters listed in Appendix 1-table 4. We simulated a network of $N$ inhibitory neurons *(threshold_lin_rate_ipn,* (*75*)), which follow the dynamical equation

$$\tau \frac{\mathrm{d}z_i}{\mathrm{d}t} = -z_i + \sum_j J_{ij}\nu_j + \mu_{\mathrm{ext},i} + \xi_i \sqrt{\tau}\sigma_{\mathrm{noise},i} \quad , \tag{55}$$

where $z_i$ is the input to neuron $i$, $\nu$ the output firing rate with (threshold linear activation function)

$$\nu = \phi(z) = \begin{cases} 0 & \text{for } z \leq 0 \\ z & \text{for } z > 0 \end{cases} \quad ,$$

time constant $\tau$, connectivity matrix $\boldsymbol{J}$, a constant external input $\mu_{\text{ext},i}$, and uncorrelated Gaussian white noise $\langle \xi_i(t) \rangle = 0$, $\langle \xi_i(s)\xi_j(t) \rangle = \delta_{ij}\delta(s-t)$, with noise strength $\sqrt{\tau}\sigma_{\text{noise},i}$. The neurons were connected using the *fixed_indegree* connection rule, with connection probability $p$, indegree $K = p \cdot N$, and delta-synapses *(rate_connection_instantaneous)* of weight $w$.

The constant external input $\mu_{\text{ext},i}$ to each neuron was normally distributed, with mean $\mu_{\text{ext}}$, and standard deviation $\sigma_{\text{ext}}$. It was used to set the firing rates of neurons, which, via the effective connectivity, influence the intrinsically generated covariances in the network. The two parameters $\mu_{\text{ext}}$ and $\sigma_{\text{ext}}$ were chosen such that, in the stationary state, half of the neurons were expected to be above threshold. Which neurons are active depends on the realization of $\mu_{\text{ext},i}$ and is therefore different for different networks.

To assess the distribution of firing rates, we first considered the static variability of the network and studied the stationary solution of the noise-averaged input $\langle z \rangle_{\text{noise}}$, which follows from Equation (55) as

$$\langle z_i \rangle_{\text{noise}} = \sum_j J_{ij} \langle \nu_j \rangle_{\text{noise}} + \mu_{\text{ext},i}. \tag{56}$$

Note that $\langle \nu_j \rangle_{\text{noise}} = \langle \phi(z_j) \rangle_{\text{noise}}$, through the nonlinearity $\phi$, in principle depends on fluctuations of the system. This dependence is, however, small for the chosen threshold linear $\phi$, which is only nonlinear in the point $z = 0$.

The derivation of $\mu_{\text{ext}}$ is based on the following mean-field considerations: according to Equation (56) the mean input to a neuron in the network is given by the sum of external input and recurrent input

$$\mu = \mu_{\text{ext}} + \mu_{\text{recurrent}} = \mu_{\text{ext}} + Kw\text{Mean}(\nu) \quad .$$

The variance of the input is given by

$$\sigma^2 = \sigma^2_{\text{ext}} + \sigma^2_{\text{recurrent}} = \sigma^2_{\text{ext}} + Kw^2\text{Var}(\nu) \quad .$$

The mean firing rate can be calculated using the diffusion approximation (*76, 77*), which is assuming a normal distribution of inputs due to the central-limit theorem, and the fact that a linear threshold neuron only fires if its input is positive

$$\begin{aligned} \text{Mean}(\nu) &= \int_{-\infty}^{\infty} \mathrm{d}\nu \, \mathcal{P}\left(\mu, \sigma^2, \nu\right) \nu \\ &= \int_{-\infty}^{\infty} \mathrm{d}z \, \mathcal{N}\left(\mu, \sigma^2, z\right) \phi(z) \\ &= \int_{0}^{\infty} \mathrm{d}z \, \mathcal{N}\left(\mu, \sigma^2, z\right) z \\ &= \frac{\sigma}{\sqrt{2\pi}} \exp\left(-\frac{\mu^2}{2\sigma^2}\right) + \frac{\mu}{2}\left[1 + \text{erf}\left(\frac{\mu}{\sqrt{2}\sigma}\right)\right] \quad , \end{aligned}$$

where $\mathcal{P}$ denotes the probability density of the firing rate $\nu$. The variance of the firing rates is given by

$$\begin{aligned} \text{Var}(\nu) &= \text{Mean}(\nu^2) - \text{Mean}(\nu)^2 \\ &= \frac{\mu^2}{4}\left[1 - \text{erf}^2\left(\frac{\mu}{\sqrt{2}\sigma}\right)\right] + \frac{\sigma^2}{2}\left[1 - \frac{1}{\pi}\exp\left(-\frac{\mu^2}{\sigma^2}\right) + \text{erf}\left(\frac{\mu}{\sqrt{2}\sigma}\right)\right] + \frac{\mu\sigma}{\sqrt{2\pi}}\text{erf}\left(\frac{\mu}{\sqrt{2}\sigma}\right) \quad . \end{aligned}$$

The number of active neurons is the number of neurons with a positive input, which we set to be equal to $N/2$

$$\begin{aligned} \frac{N}{2} &\overset{!}{=} N \int_0^\infty \mathrm{d}z \, \mathcal{N}\left(\mu, \sigma^2, z\right) \\ &= \frac{N}{2}\left[1 + \mathrm{erf}\left(\frac{\mu}{\sqrt{2}\sigma}\right)\right] \quad , \end{aligned}$$

which is only fulfilled for $\mu = 0$. Inserting this condition simplifies the equations above and leads to

$$\mu_{\mathrm{ext}} = -\frac{Kw\sigma}{\sqrt{2\pi}} \quad .$$

For the purpose of relating synaptic weight $w$ and spectral bound $R$, we can view the nonlinear network as an effective linear network with half the population size (only the active neurons). In the latter case, we obtain

$$w = -\frac{R}{\sqrt{\frac{N}{2}p(1-p)}} \quad .$$

For a given spectral bound $R$, this relation allows us to derive the value

$$\mu_{\mathrm{ext}} = \sqrt{\frac{Np}{\pi(1-p) - (\pi - 1)R^2}} \sqrt{\sigma_{\mathrm{ext}}^2} R \quad , \tag{57}$$

that, for a arbitrarily fixed $\sigma_{\mathrm{ext}}$ (here $\sigma_{\mathrm{ext}} = 1$), makes half of the population being active. We were aiming for an effective connectivity with only weak fluctuations in the stationary state. Therefore, we fixed the noise strength for all neurons to the small value $\sigma_{\mathrm{noise}} = 0.1 \ll \sigma_{\mathrm{ext}}$ compared to the external input, such that the noise fluctuations did not have a large influence on the calculation above that determines which neurons were active.

To show the effect of a change in the effective connectivity on the covariances, we simulated two networks with identical connectivity, but supplied them with slightly different external inputs. This was realized by choosing

$$\mu_{\mathrm{ext},i}^{(\alpha)} = \mu_{\mathrm{ext},i} + \mu_{\mathrm{ext},i}^{(\alpha)} \quad ,$$

with

$$\begin{aligned} \mu_{\mathrm{ext},i} &\sim \mathcal{N}\left(\mu_{\mathrm{ext}}, [1-\epsilon]\, \sigma_{\mathrm{ext}}^2\right) \quad , \\ \mu_{\mathrm{ext},i}^{(\alpha)} &\sim \mathcal{N}\left(0, \epsilon\sigma_{\mathrm{ext}}^2\right) \quad , \end{aligned}$$

$\epsilon \ll 1$, and $\alpha \in \{1, 2\}$ indexing the two networks. The main component $\mu_{\mathrm{ext},i}$ of the external input was the same for both networks. But, the small component $\mu_{\mathrm{ext},i}^{(\alpha)}$ was drawn independently for the two networks. This choice ensures that the two networks have a similar external input distribution (figure 5$B$1 of the main text), but with the external inputs distributed differently across the single neurons (figure 5$B$2 of the main text). How similar the external inputs are distributed across the single neurons is determined by $\epsilon$.

The two networks have a very similar firing rate distribution (figure 5$E$1 of the main text), but, akin to the external inputs, the way the firing rates are distributed across the single neurons differs between the two networks (figure 5$E$2 of the main text). As the effective connectivity depends on the firing rates

$$W_{ij} = J_{ij}\phi'(\nu_j) \quad ,$$

this leads to a difference in the effective connectivities of the two networks and therefore to different covariance patterns, as discussed in figure 5 of the main text.

| **Network Parameters** | | |
|---|---|---|
| $N$ | 2000 | Number of neurons |
| $p$ | 0.1 | Connection probability |
| $\tau$ | 1 ms | Time constant |
| $\sigma_\mu$ | 1 Hz | Standard deviation of external input |
| $\sigma_{\mathrm{noise}}$ | 0.1 Hz | Standard deviation of noise |
| $R$ | $[0.1, 0.2, ..., 0.9]$ | Spectral bound |
| $\epsilon$ | 0.1 | Parameter controlling difference of two network simulations |
| **Simulation Parameters** | | |
| $\mathrm{d}t$ | 0.1 ms | Simulation step size |
| $t_{\mathrm{init}}$ | 100 ms | Initialization time |
| $t_{\mathrm{sim}}$ | 2000000 ms | Simulation time without initialization time |
| $t_{\mathrm{sample}}$ | 1 ms | Sample resolution at which rates where recorded |
| **Analysis Parameters** | | |
| $N_{\mathrm{sample}}$ | 200 | Sample size |
| $T$ | 100 ms | Correlation time window |

Appendix 1-table 4: Parameters used for NEST simulation and subsequent analysis.

We performed the simulation for spectral bounds ranging from $0.1$ to $0.9$ in increments of $0.1$. We calculated the correlation coefficient of firing rates and the correlation coefficient of time-lag integrated covariances between $N_{\mathrm{sample}}$ neurons in the two networks (figure 5*D* of the main text) and studied the dependence on the spectral bound.

To check whether the simulation was long enough to yield a reliable estimate of the rates and covariances, we split each simulation into two halves, and calculate the correlation coefficient between the rates and covariances from the first half of the simulation with the rates and covariances from the second half. They were almost perfectly correlated (figure 5*C* of the main text). Then, we calculated the correlation coefficients comparing all halves of the first simulation with all halves of the second simulation, showing that the covariance patterns changed much more than the rate patterns (figure 5*C* of the main text).
